# Astrocytes and neurons share brain region-specific transcriptional signatures

**DOI:** 10.1101/2020.04.21.038737

**Authors:** Álvaro Herrero-Navarro, Lorenzo Puche-Aroca, Verónica Moreno-Juan, Alejandro Sempere-Ferràndez, Ana Espinosa, Rafael Susín, Laia Torres-Masjoan, Eduardo Leyva-Díaz, Marisa Karow, María Figueres-Oñate, Laura López-Mascaraque, José P. López-Atalaya, Benedikt Berninger, Guillermina López-Bendito

**Affiliations:** Instituto de Neurociencias de Alicante, Universidad Miguel Hernández-Consejo Superior de Investigaciones Científicas (UMH-CSIC), Sant Joan d’Alacant, Spain; Neural Coding Laboratory, Istituto Italiano di Tecnologia, Via Morego 30, 16163 Genova, Italy; Department of Biological Sciences, Howard Hughes Medical Institute, Columbia University, New York, NY 10027, USA; Institute of Psychiatry, Psychology, and Neuroscience, Centre for Developmental Neurobiology, and MRC Centre for Neurodevelopmental Disorders, King’s College London, SE1 1UL London, UK; Institute of Biochemistry, Friedrich-Alexander-University Erlangen-Nürnberg, 91054 Erlangen, Germany; Physiological Genomics, Biomedical Center, Ludwig Maximilians University Munich, 82152 Planegg/Munich, Germany; Instituto Cajal, CSIC, Madrid, Spain; Max Planck Research Unit for Neurogenetics, Max von-Laue-Strasse 4, 60438 Frankfurt, Germany; Institute of Physiological Chemistry, University Medical Center Johannes Gutenberg University Mainz, 55128 Mainz, Germany

**Author notes:** These authors contributed equally to this work (José P. López-Atalaya, Benedikt Berninger).

## Abstract

Neuronal cell diversity is essential to endow distinct brain regions with specific functions. During development, progenitors within these regions are characterised by specific gene expression programs, contributing to the generation of diversity in postmitotic neurons and glia. While the region-specific molecular diversity of neurons and astrocytes is increasingly understood, whether these cells share region-specific programs remains unknown. Here, we show that in the neocortex and thalamus, neurons and astrocytes express shared region-specific transcriptional and epigenetic signatures. These signatures not only distinguish cells across brain regions but are also detected across substructures within regions, such as distinct thalamic nuclei, where clonal analysis revealed the existence of common nucleus-specific progenitors for neurons and glia. Consistent with their shared molecular signature, regional specificity was maintained following astrocyte-to-neuron reprogramming. A detailed understanding of these regional-specific signatures may thus inform strategies for future cell-based brain repair.

---

The development of neuronal diversity is central for the organization and function of the CNS. This diversity is largely determined by specific transcriptional programs already expressed at the progenitor stage^1-7^. These programs can undergo temporal regulation allowing for sequential generation of different progeny from the same original progenitor^4,8^. The most drastic case of this temporal regulation occurs at the switch of progenitors from neurogenic to gliogenic competence^9^. Moreover, transcriptional programs are also diversified across brain regions to define the positional identity of the progenitors. Pioneering work in the spinal cord suggests that the diversification of astrocytes might follow the same organizing principle of positional identity^10^. This notion has recently received further support by clonal analyses and single cell transcriptomics that unveiled highly characteristic distributions within brain regions of heterogeneous glia^11-14^. However, given that neurons and astroglia are generated from the same germinal zones, they could share common molecular signatures reflecting their origin and potentially acting to coordinate region-specific developmental features. Here, we address this possibility and report that thalamic and cortical astrocytes have region-specific transcriptional and epigenetic signatures, which are shared with the neurons generated within the same thalamic or cortical progenitor domain but not beyond. These shared signatures confer a remarkable degree of regional specification for astrocyte-to-neuron reprogramming induced by the proneural factor Neurogenin 2. Finally, manipulating the regional-specific code in defined astrocyte populations redirects reprogramming toward neurons of different, yet predictable regional identity.

## Shared gene expression signatures between astroglia and neurons

To test the hypothesis that astrocytes and neurons generated within the same brain region share molecular signatures unique to this region, we identified differentially expressed genes (DEGs) between astrocytes of the thalamus and cortex, performed a similar analysis between thalamic and cortical neurons, and then searched for potential overlap among the two sets of DEGs (Fig. **1a**). We performed bulk RNA-sequencing (RNA-seq) on astrocytes isolated from thalamus (comprising dorsolateral geniculate dLG, ventral posteromedial VPM and ventro-medial geniculate MGv nuclei) and primary somatosensory cortex (S1) using astrocyte reporter mice (*Gfap::Gfp*) at postnatal day 7 (P7) (Extended data Fig. **1a, b**), well-after the peak of astrogenesis^15^. Principal component analysis (PCA) revealed that thalamic and cortical astrocytes clustered according to their anatomical origins (Fig. **1b**). This analysis revealed 1675 versus 1287 DEGs enriched in thalamus and cortex, respectively (Fig. **1c** and Extended Fig. **1c**). Among the DEGs enriched in each population, we identified several genes, including transcription factors, that are known to be highly expressed in neurons of the respective regions^16,17^ (Fig. **1c** and Extended Fig. **1d**). This prompted us to perform gene ontology (GO) over-representation and gene set enrichment analyses (GSEA) of the DEGs between thalamic and cortical astrocytes, which revealed marked differences in developmental programs and distinct region-specific molecular pathways that have been previously associated with neurons from these regions (Fig. **1d** and Extended Data Fig. **1e, f**). To unveil region-specific genes shared among astrocytes and neurons of the corresponding regions, we first identified the most highly DEGs enriched in thalamic and cortical neurons, by comparing RNA-seq data of neurons isolated at P0 from a thalamic reporter line (*Gbx2-Cre::Tomato-floxed*) with a published dataset of P1 cortical neurons^17^ (Extended Data Fig. **2a-c**). We found that genes specifically enriched in thalamic or cortical neurons were significantly overrepresented among DEGs in thalamic or cortical astrocytes, respectively. Among the 400 most differentially expressed genes in thalamic neurons, only 6% were shared with cortical astrocytes, whereas 32.75% of these genes were significantly expressed by thalamic astrocytes, albeit at a lower level (including *Lef1, Gbx2* or *Rorα*; Fig. **1e, f** and Extended Data Fig. **2d-g** and **3**). A significant overlap in gene expression, 17.5%, was also observed for cortical neurons and astrocytes, where genes like *Tbr1* or *Fezf2* were identified in both populations. Next, we interrogated the overlap in expression of region-specific genes between neurons and astrocytes at single cell level by analyzing an independent, published data set containing single cell transcriptomes of thalamic and cortical neurons and astrocytes from adult mice^18^ (Extended Data Fig. **4**). Despite the low capture efficiency of the scRNA-seq, this analysis confirmed our result that astrocytes and neurons of thalamic and cortical origin share region-specific gene expression programs, and that those similarities are conserved also in the adult.

**Fig. 1.**
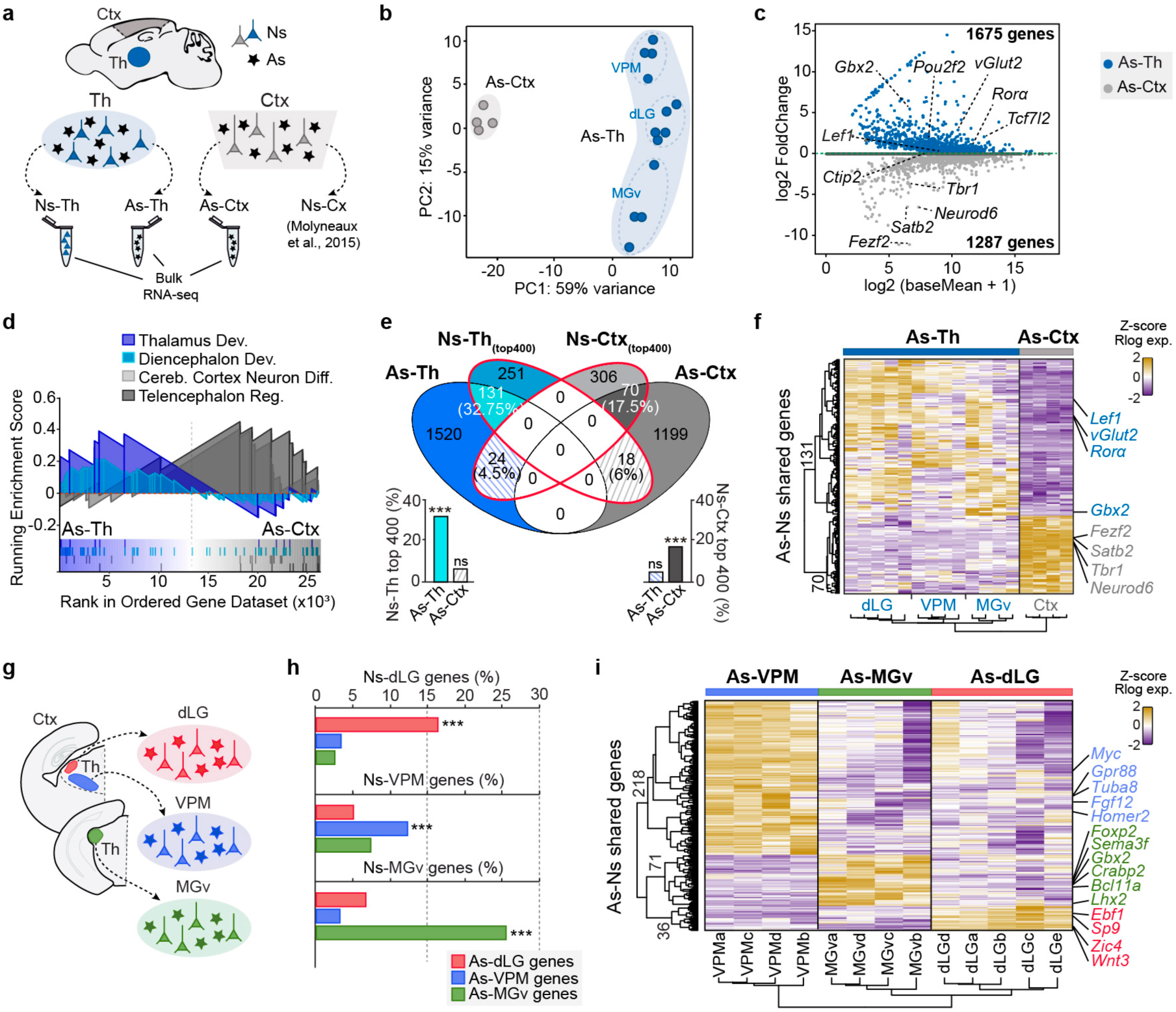
Astrocytes and neurons share region-specific transcriptome profiles. **a**, Schematic of the RNA-seq experiments that include publicly available data for cortical neurons^17^. **b**, Principal Component Analysis (PCA) of the transcriptomes of astrocytes (As) from thalamus (Th), including dLG (*n* = 5 samples), VPM (*n* = 4) and MGv (*n* = 4), and cortex (Ctx, *n*= 4) at P7. **c**, MA plot displaying differentially expressed genes (DEGs). Blue and light grey dots represent thalamic and cortical DEGs with their mean normalized counts, respectively. Dark grey dots represent genes that failed to give a significant result. **d**, Enrichment plots from GSEA of two specific GO terms related to the thalamic and cortical formation. **e**, Venn diagram showing the genes that overlap between As and neurons (Ns) in both the thalamus and cortex. Ns-Th, included dLG (*n* = 4), VPM (*n* = 4) and MGv (*n* = 3). Ns-Ctx included Callosal Projecting Neurons (CPN, *n* = 2); Corticothalamic projecting neurons (CThPN, *n* = 2); and Subcerebral projecting neurons (ScPN, *n* = 2). Bar plot represents the proportion of shared genes. **f**, Heatmap showing overlapping genes between As and Ns in the thalamus and cortex. **g**, Schematic showing the bulk RNA-seq for As and Ns in specific thalamic nuclei. **h**, Bar plots represent the percentage of gene overlap between As and Ns from each thalamic nucleus. **i**, Heatmap showing the overlapping DEGs between As and Ns in each nucleus. Each column represents a biological replicate and the color-code represents the Z-score normalized expression (upregulated genes in yellow, downregulated genes in purple).

We next asked whether region-specific gene expression programs can be identified at the level of individual regional subdivisions such as those of sensory thalamic nuclei. Thus, we compared the transcriptomes of astrocytes and neurons from the three main sensory thalamic nuclei (dLG, VPM, and MGv). PCA analysis identified three well-defined clusters corresponding to each nucleus in both astrocytes and neurons, supporting the notion that the identity of each thalamic nucleus is encoded transcriptionally in a cell-type independent manner (Extended Data Fig. **5a-d**). Hence, the nucleus-specific DEGs of astrocytes exhibited a significant overlap with those of the neurons from the same nucleus (e.g., *Sp9* for the dLG or *Crabp2* for MGv: Fig. **1g-i** and Extended Data Fig. **5e**), although the expression levels of these genes were significantly lower in the astrocytic populations (Extended Data Fig. **5f**). In summary, our results demonstrate that region- and subregion-specific transcriptional programs are shared by astrocytes and neurons.

## Thalamic progenitor clones are nucleus specific

Next, we investigated whether the significant gene expression overlap between postmitotic astrocytes and neurons reflects a common clonal origin during embryonic development. This would imply that within the thalamus, cells belonging to the same clone should not disperse beyond nuclear boundaries. To test this hypothesis, we first analysed the distribution of astrocytes originating from single clones across thalamic sensory nuclei. We tracked astrocyte clones arising from E11.5 progenitors by electroporating a battery of plasmids encoding distinct fluorophores under the control of the *Gfap* promoter, following transposase mediated integration (“StarTrack”)^11^, and analyzed the dipersion of each clone at P8 (Fig. **2a, b** and Extended Data Fig. **6a, b**). This revealed that clonally related astrocytes remain within the boundaries of a given nucleus with little dispersion to other nuclei, even in the case of larger clones (>10 cells) (Fig. **2c** and Extended Data Fig.**6c**), extending and confirming previous studies^19^. Next, we addressed the question of whether thalamic progenitors that generate astrocytes also produce neurons, and if so, whether these neurons stay within the same nuclear boundaries as their sibling astrocytes. Thalamic clones containing neuronal and/or non-neuronal cells were tracked by using the same set of fluorophores under the control of an ubiquitously expressed promoter^20^ (Fig. **2d, e** and Extended Data Fig. **6d**). While we found 39% of clones consisting only of neurons or glia, the majority (61%) were mixed, containing similar proportions of neurons and glia (Fig. **2f** and Extended Data Fig. **6e**). We found that mixed clones covered territories that largely respected nuclear boundaries, although neurons exhibited a wider range of dispersion (Fig. **2g, h** and Extended Data Fig. **6f**). Our data suggest that the overlap in region-specific gene expression between neurons and glia of each sensory thalamic nucleus is the result of their common clonal origin together with the limited spatial dispersion of clonally-related cells, and may indicate that positional information is retained from an early progenitor stage on.

**Fig. 2.**
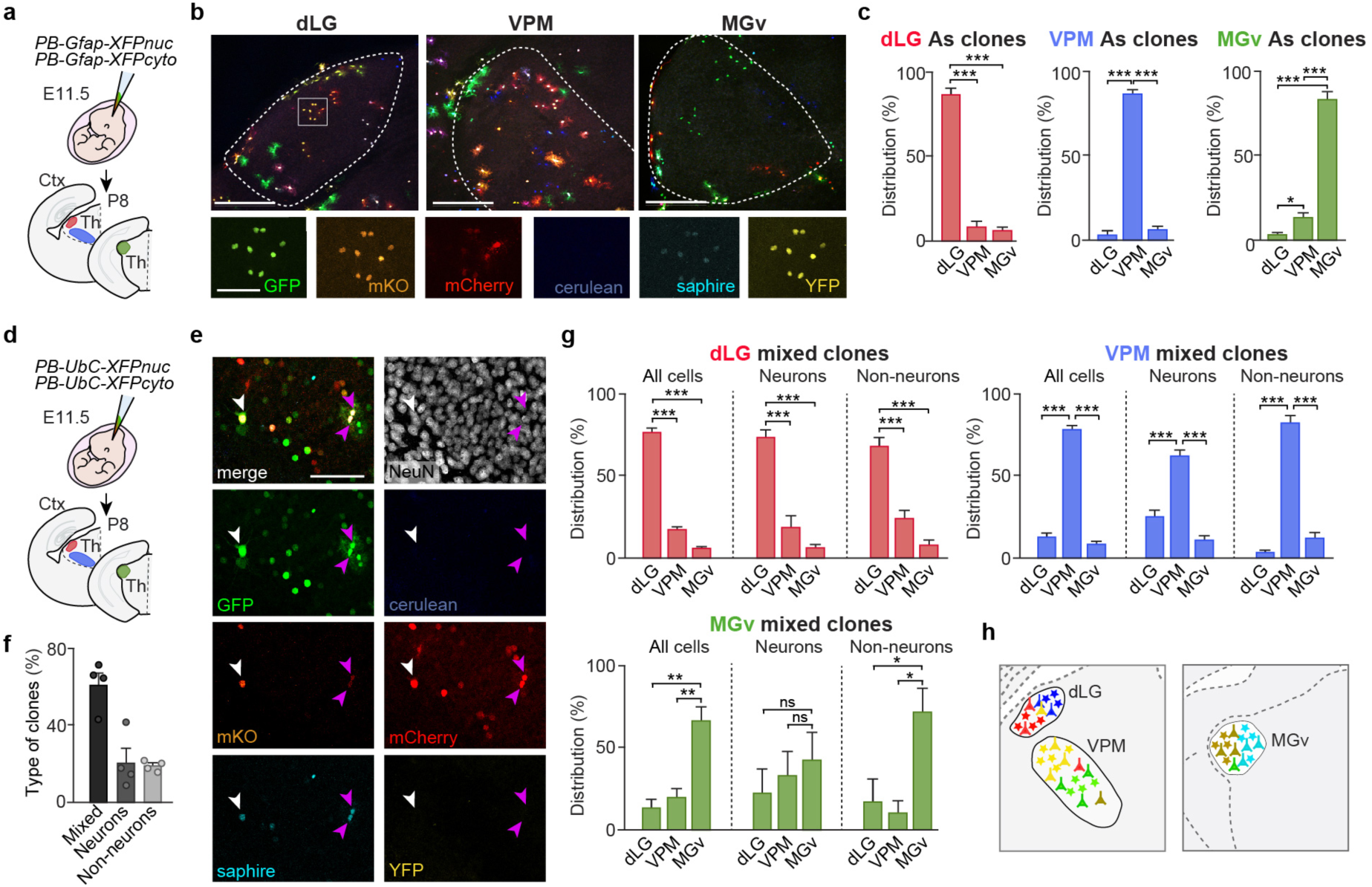
Clonally related astrocytes and neurons remain within the same nuclear boundaries. **a**, Experimental design for the analysis of astrocytic clones in the sensory nuclei (dLG, VPM, MGv). **b**, Thalamic astrocytes labelled with the GFAP*-StarTrack* constructs at P8. Insets show the expression of each fluorescent reporter in a dLG astrocyte clone (white square). **c**, Quantification of the dispersion of the clonally related astrocytes (*n* = 320 clones from 5 electroporated animals). **d**, Experimental design for the study of neuronal and non-neuronal clonal cells with the UbC*-StarTrack* constructs. **e**, Example of a neuron (white arrow) and two astrocytes (purple arrows) from the VPM coming from the same progenitor, thus sharing the same color code. **f**, Type of analyzed clones based on their cell type composition (*n* = 4 electroporated animals). **g**, Quantification of the dispersion of clonally related neuronal and non-neuronal cells from mixed clones, in the different thalamic sensory nuclei (*n* = 130 clones from 4 electroporated animals). **h**, Schema representing the specificity in the nuclei-dependent localization of clonal cells. Data are the means ± s.e.m. Scale bars, 100 μm.

## Astrocytes reprogram into region specific neurons

Since forebrain astrocytes and neurons share region-specific gene expression, we hypothesized that such molecular signature could instruct transcription factor-induced reprogramming of astrocytes towards an identity akin to their sibling neurons. To test this hypothesis, we injected a retrovirus encoding the proneural gene *Neurogenin 2* (*Neurog2*) and the cell-death regulator Bcl2^21^ into the somatosensory cortex and thalamus of P3 mice (Fig. **3a**). At this developmental stage, retroviruses only transduce proliferative glia^15^. Transduction with a retrovirus encoding *Bcl2* and *Gfp* alone, as control, resulted in labelling of glial cells (Extended Data Fig.**7a**). In contrast, transduction with *Neurog2 Bcl2* encoding retrovirus resulted in the appearance of numerous induced neurons (Extended Data Fig.**7a, b**). Consistent with our hypothesis, *in vivo* induced neurons expressed markers specific for a thalamic (*Lef1* and *Rorα*) or cortical (*Tbr1* and *Ctip2*) neuronal identity despite the fact that they were induced with the same transcription factor (Fig. **3b** and **c**).

**Fig. 3.**
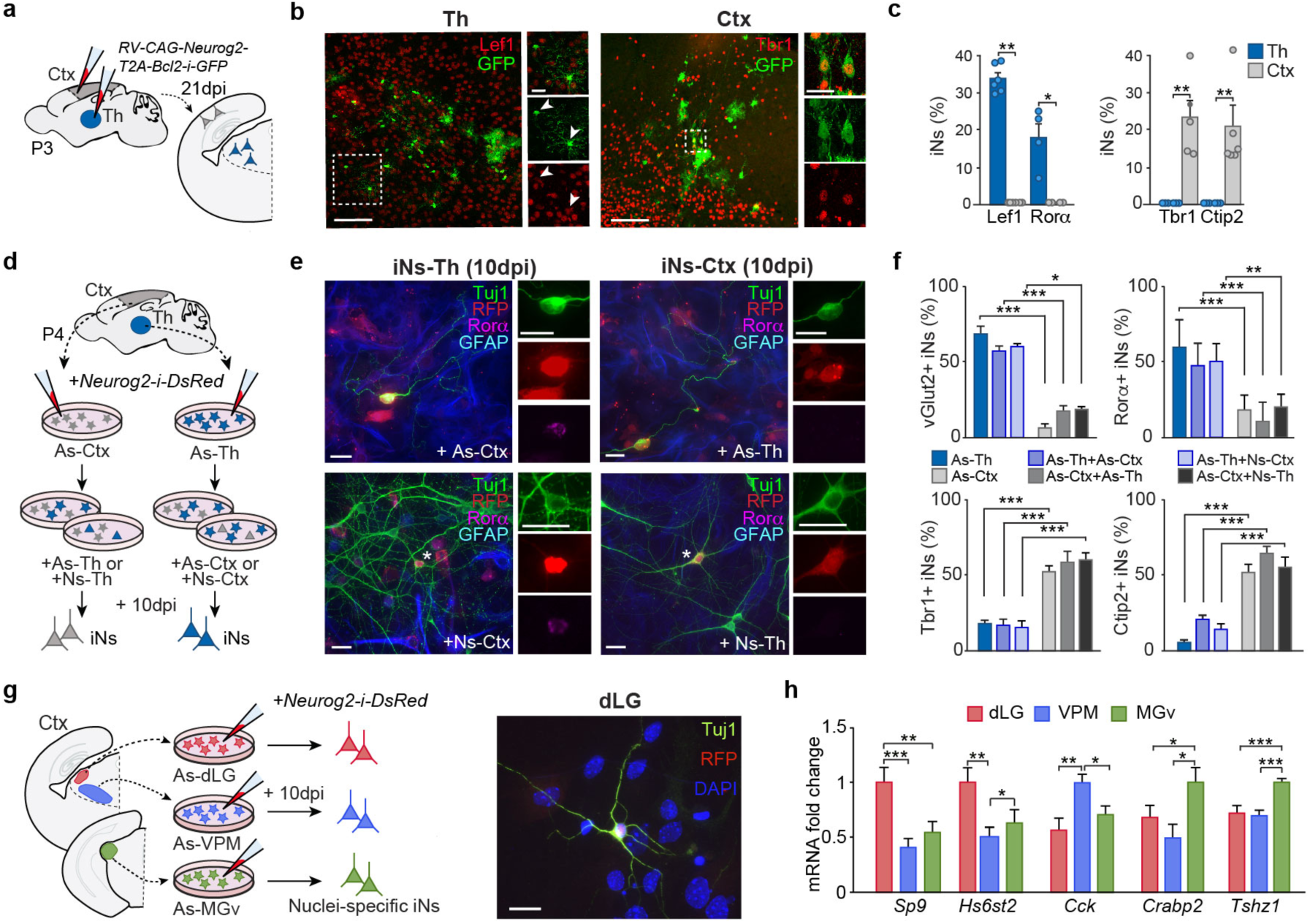
Astrocytes are reprogrammed into region-specific neurons. **a**, Experimental design for the *in vivo* reprogramming. **b**, Immunofluorescence for thalamic and cortical markers in iNs reprogrammed from cortical or thalamic astrocytes *in vivo*. **c**, Percentage of iNs expressing thalamic or cortical markers after reprogramming *in vivo* (*n* = 4-6 injected mice). **d**, Experimental design for assessing the influence of the environment on the induced neurons identity. **e**, Immunostaining for the thalamic marker Rorα in cortical or thalamic iNs (RFP+/Tuj1+) in the different conditions. **f**, Quantification of the percentage of iNs generated from cortical or thalamic astrocytes that express *vGlut2, Rorα, Tbr1* or *Ctip2* in control conditions or when mixed with astrocytes or neurons from the thalamus or the cortex, respectively (*n* = 6-14 independent cultures per condition). **g**, Left, experimental design. Right, image of an iN from dLG astrocytes at 10 days post infection (dpi). **h**, RT-qPCR showing the expression of specific neuronal genes in the iNs after 10 dpi (*n* = 10-14 independent cultures per condition). Data are means ± s.e.m. Dots in **c** represent every single value. Scale bars, 100μm in **b** (insets 25μm), and 25μm in **e** and **g**.

Our data suggests that reprogramming of astrocytes into region-specific neurons is a consequence of their shared gene expression through a common lineage. However, it does not exclude the possibility that region-specific reprogramming is influenced by environmental signals provided by other local cells. To test this, we cultured astrocytes from thalamus and cortex and examined their newly acquired neuronal identity for region specific gene expression following reprogramming by *Neurog2* (Extended Data Fig.**7c, d)**. As observed *in vivo*, thalamic and cortical induced neurons exhibited signatures of thalamus and cortex, respectively, as shown by the differential expression of thalamic markers such as *vGlut2, Rorα, Gbx2, Pou2f2* or *Lef1*, or cortical markers such as *Tbr1* or *Ctip2* (Extended Data Fig. **8**). To exclude a significant role of the environment in specifying the regional identity of induced neurons, we co-cultured thalamic or cortical astrocytes undergoing reprogramming with neurons or astrocytes from the cortex or thalamus, respectively. Neurons induced from thalamic astrocytes expressed thalamic markers, irrespective of whether they had been cultured alone or with cortical cells. Conversely, cortical astrocytes gave rise to neurons expressing cortical markers irrespective of the culture conditions (Fig. **3d-f**). These experiments revealed that the regional identity of induced neurons is largely cell autonomous.

Finally, as astrocytes and neurons from distinct thalamic sensory nuclei share expression of nucleus-specific genes, we hypothesized that reprogramming of thalamic astrocytes may yield neurons with nucleus specific signatures. To this end, we isolated and reprogrammed astrocytes from dLG, VPM and MGv *in vitro* with Neurog2 (Fig. **3g**). We found that induced neurons derived from dLG astrocytes expressed dLG-specific genes *Sp9, Hs6st2*, while those derived from MGv astrocytes expressed MGv-specific genes *Crabp2* and *Tshz1*. Lastly, induced neurons of VPM astrocyte origin expressed the VPM-marker *Cck* (Fig. **3h**)^16^. Taken together, these results show that Neurog2 triggers specific neuronal gene expression in astrocytes that reflects their place of origin.

## *Gbx2* re-specifies cortical astrocytes towards thalamic fate

The aforementioned results strongly suggest that transcriptional signatures shared between neurons and astrocytes drive regional specification of the latter during neuronal reprogramming. To directly test this, we examined whether co-expression of a thalamic fate determinant *Gbx2*^22^, a factor being shared between astrocytes and neurons of the thalamus, could re-direct neuronal reprogramming of cortical astrocytes towards a thalamic identity (Fig. **4a**). Whereas in cortical astrocytes Neurog2 induced the expression of the cortical neuron fate determinants *Tbr1* and *Ctip2*, co-expression of *Gbx2* strongly suppressed this. Moreover, combined expression of *Neurog2* and *Gbx2* increased thalamic signature genes like *Pou2f2* and v*Glut2* in cortical astrocytes (Fig. **4b**). This data provides strong support for partial re-direction of neuronal reprogramming towards a thalamic identity (Fig. **4c**). In thalamic astrocytes, by contrast, *Neurog2* sufficed for inducing significant expression of *Pou2f2* and *vGlut2* expression (Fig. **4b**). Genes that displayed differential regulation by *Neurog2* with or without *Gbx2*, exhibited an epigenetically poised state in cortical or thalamic astrocytes, as determined by the ratio of active (H3K4me3) and repressive (H3K27me3) histone marks in their proximal regulatory elements (Fig. **4d**, Extended Data Fig. **9** and **10a-b**). In contrast, non-responsive genes (*Fezf2, Rorα, Lef1*) exhibited significant origin-dependent baseline expression both transcriptionally and epigenetically in thalamic and cortical astrocytes (Fig. **4b, d**, Extended Data Fig. **9a-c** and **10a-b**).

**Fig. 4.**
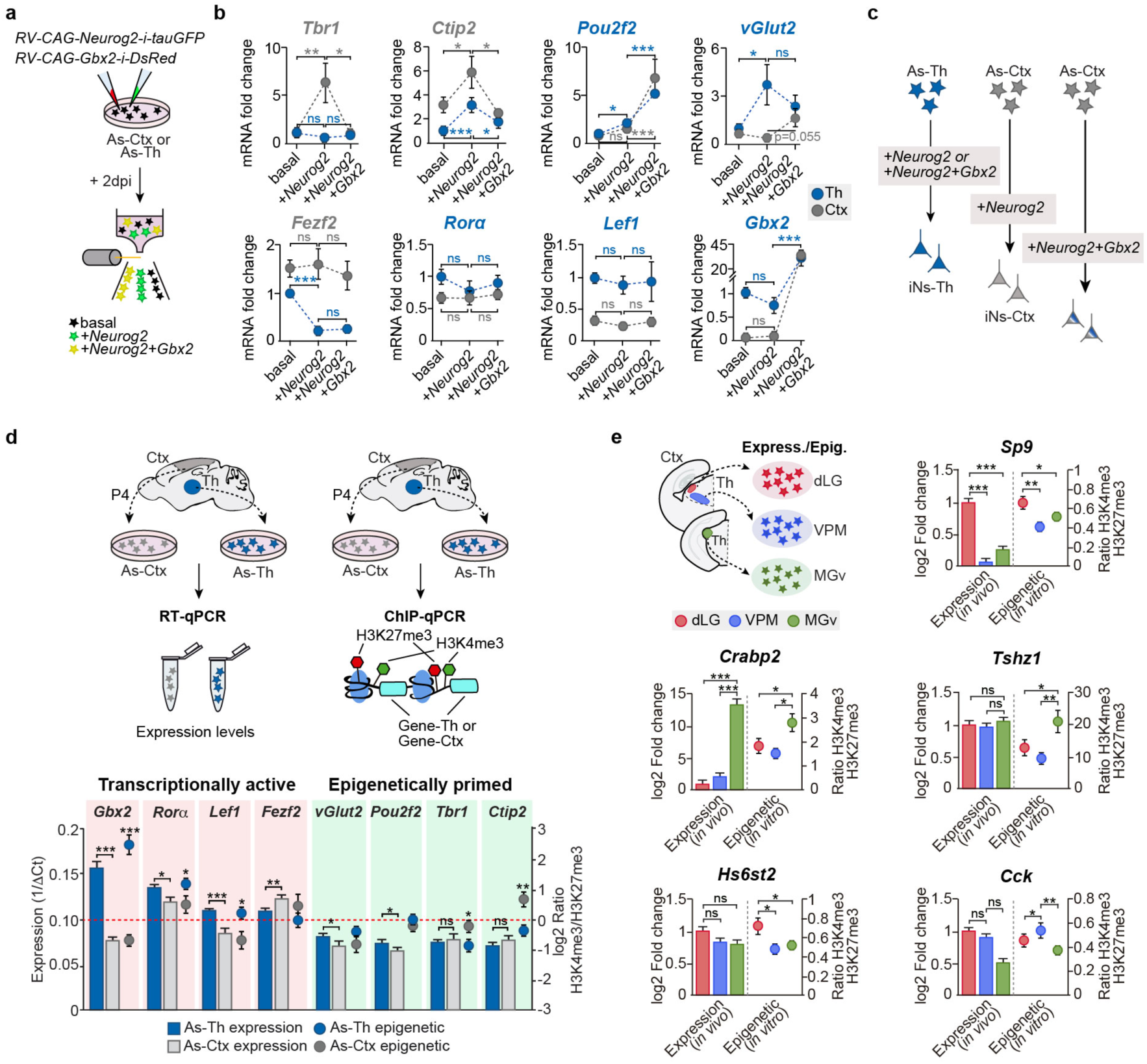
Poised epigenetic state of region-specific gene expression in astrocytes. **a**, Experimental design. **b**, Quantification of specific genes expression by RT-qPCR in astrocytes in basal conditions and after the overexpression of *Neurog2* alone, or with a thalamic specific gene (*Gbx2*) (*n* = 6-14 independent cultures per condition). **c**, Schematic conclusion of the experiment. **d**, Bars representing the average basal expression levels of the studied genes in thalamic and cortical astrocytes (right axis), and dots showing the epigenetic state of the promoter of those genes, in terms of the presence of two histone marks, H3K4me3 and H3K27me3 (left axis) (*n* = 12-23 independent ChIP samples per condition). Red dashed line indicates the point where H3K4me3 and H3K27me3 marks are present at the same level. **e**, Graphs showing the H3K4me3 and H3K27me3 ratio *in vitro* (*n* = 14-18 independent ChIP samples per condition), and the basal *in vivo* expression of neuronal specific genes in thalamic astrocytes from each nucleus. Data are means ± s.e.m.

Finally, we addressed the question of whether a similar epigenetically poised state might explain the differential induction of nuclei-specific neuronal genes in astrocytes of distinct thalamic territories (Fig. **3g, h**). To this end, we first compared basal expression levels and presence of active (H3K4me3) and repressive (H3K27me3) epigenetic marks at proximal regulatory elements of these genes, known to be differentially expressed in dLG, VPM and MGv neurons (Extended Data Fig. **5**, and ^16^). Intriguingly, irrespective of the baseline expression level, these genes exhibited an active (*Sp9, Crabp2, Tshz1*) or poised/less repressed (*Hs6st2, Cck*) epigenetic state of their proximal regulatory elements consistent with their nuclear origin (Fig. **4e** and Extended Data Fig. **9d** and **10c, d**). Nucleus-specific epigenetic priming might explain the observed differential transcriptional responsiveness to Neurog2 of genes whose levels of transcription are indistinguishable across nuclei prior to reprogramming.

## DISCUSSION

Using genome wide analysis, we show that astrocytes of different brain regions actively transcribe genes that also correspond to regional genes in neurons. This remarkable relatedness between astrocytes and neurons from the same brain region correlates with their shared clonal origin, as shown for distinct sensory nuclei of the thalamus. Furthermore, region-specific molecular signatures create a strong bias intrinsic to astrocytes towards generating neurons of appropriate regional identity when reprogrammed by the proneural factor Neurog2. Thus, intrinsic differences between astrocyte populations may facilitate strategies to repair diseased neural circuits in a region-specific manner.

Despite their common developmental origin, neurons and astrocytes constitute cell types easily distinguishable by their morphological and electrophysiological properties. However, our study reveals that these two cell types show an unexpected overlap in the expression of genes that confer regional identity. Such overlap can be found at single cell level and extends into adulthood. Among the shared genes, there was a significant amount of transcription factors, many of which play well-described roles in neuron subtype specification (e.g. *Gbx2, Lef, Fefz2, Tbr1*)^22-25^. What might be the function of sharing these region-specific genes between neurons and astrocytes? We envisage three scenarios: Firstly, shared gene expression may be a vestige of shared clonal origin without any specific function beyond development; secondly, these genes may adopt different functions in astrocytes as compared to neurons; finally, expression of these genes might act as a code to facilitate region-specific interactions of astrocytes with their sibling neurons. While these scenarios are not mutually exclusive, the latter may provide an attractive mechanism by which neurons could modulate the spatial distribution of astrocytes^12^. Indeed, our clonal analyses reveals that neurons and sibling astrocytes originating from the same thalamic progenitor clone populate very similar territories, respecting boundaries among thalamic nuclei, extending earlier observations of the existence of nucleus-specific progenitor domains in the thalamus^19,26^.

It seems plausible that the shared region-specific gene expression is accounted for by epigenetic signatures inherited from a common progenitor and maintained throughout postmitotic development. Indeed, our data provide evidence for region-specific differences in the epigenetic state of regulatory elements of these genes in cortical and thalamic astrocytes, even up to the level of thalamic nuclear divisions. Conversely, these region-specific genes apparently escape the long-term epigenetic repression that occurs at neuronal gene loci at the developmental switch from neurogenesis to gliogenesis^27,28^. The epigenetic configuration at region-specific genes might function as a latent mechanism to keep some neuronal expressed genes in a “poised state” in astrocytes, which may become activated by reprogramming factors such as shown here. The fact that epigenetic configurations are heritable through cell divisions^4,28^ might confer astrocytes with a specific and long-lasting regional differentiation potential as may occur during injury-induced neurogenesis^29,30^. Finally, the fine-grained heterogeneity of astrocytes between and within brain regions (^10,12^ and this study) may provide a basis for reconstructing diseased brain circuits that require the generation of multiple neuron types^31,32^, with a minimal number of molecular manipulations.

## Acknowledgments

The authors are grateful to L.M. Rodríguez and B. Andrés for their excellent technical support; to V. Borrell for providing the plasmids for retroviral production; and to members of G. López-Bendito’s laboratory for stimulating discussions. We thank D. Jabaudon, A. Nieto and S. Garel for their input into the manuscript.

## Author contributions

A.H-N. and G.L-B. designed the experiments. A.H-N. conducted the astrocyte reprogramming experiments *in vitro* and *in vivo*, and the epigenetic analyses. L.P-A and J.L-A. conducted all bioinformatic analysis of massively parallel sequencing data. V.M-J. and A.H-N. conducted the *in vivo* clonal studies. A.S-F. performed the electrophysiological recordings. A.H-N., V.M-J. and A.E. collected tissue for RNAseq. R.S. produced the viruses. L.T-M. cloned the Bcl2-Neurog2 plasmid for *in vivo* reprogramming. E.L-D., M.K. and B.B. performed pioneer reprogramming experiments *in vitro* and provided the Neurog2 reprogramming construct. M.F-O. and L.L-M. provided the StarTrack constructs. G.L-B. acquired funding; A.H-N, B.B. and G.L-B. wrote the paper.

## Data and materials availability

The RNAseq data generated in this study is deposited in the NCBI Gene Expression Omnibus repository under accession GSE135602 (astrocytes of the mouse cortex and thalamic nuclei, URL: https://www.ncbi.nlm.nih.gov/geo/query/acc.cgi?acc=GSE135602) and GSE138976 (neurons of the mouse thalamic nuclei, URL: https://www.ncbi.nlm.nih.gov/geo/query/acc.cgi?acc=GSE138976). The RNAseq data generated in this study will be available upon manuscript publication. Transcriptional profiles (RNAseq) of mouse cortical neurons from Molyneaux et al.^17^ were obtained from the NCBI Gene Expression Omnibus repository (Accession GSE63482, URL: https://www.ncbi.nlm.nih.gov/geo/query/acc.cgi?acc=GSE63482). Mouse brain single-cell RNAseq data from Zeisel et al.^18^ were obtained from the NCBI Sequence Read Archive (Accession SRP135960, URL: https://www.ncbi.nlm.nih.gov/sra/SRP135960). ChIPseq coverage tracks for *In silico* Neurog2 binding sites determination were performed based on publicly available datasets from H3K4me3 (Accession experiment ENCSR258YWW, URL: https://www.encodeproject.org/experiments/ENCSR258YWW/) and H3K27me3 (Accession experiment ENCSR070MOK, URL: https://www.encodeproject.org/experiments/ENCSR070MOK/) histone marks from The ENCODE Project.

## Supplementary Materials

Extended Data Figures 1 to 10

Extended Data Tables 1 to 4

## Methods

### Mouse strains

All transgenic animals used in this study were maintained on ICR/CD-1, FVB/N-Tg or C57BL/6J genetic backgrounds and genotyped by PCR. The day of the vaginal plug was stipulated as E0.5. The *Gfap::Gfp* line (Jackson Laboratories, stock number 003257) was in a FVB/N-Tg genetic background, the *Gad67::Gfp* line^33^ in C57BL/6J, and the *R26*^*tdTomato*^ Cre-dependent line (Jackson Laboratories, Stock number 007908), and the *Gbx2*^*CreERT2/+*^ mouse^34^ lines were in a ICR/CD-1 genetic background. Tamoxifen induction of Cre recombinase in the double mutant embryos (*Gbx2*^*CreERT2/+*^::*R26*^*tdTomato*^) was performed as previously described^35^. The Committee on Animal Research at the University Miguel Hernández approved all the animal procedures, which were carried out in compliance with Spanish and European Union regulations.

### Isolation of astrocytes and neurons for RNA-seq

The brains (four brains were pooled for each sample) were extracted in ice-cold KREBS solution and cut in the vibratome in 300μm slices and cells were dissociated as in a previous publication^16^. Thalamic nuclei (dLG, VPM and MGv) and somatosensory cortex (S1) were dissected and pooled in cold dissociation medium (20 mM glucose, 0.8 mM kynurenic acid, 0.05 mM APV, 50 U/ml penicillin 0.05 mg/ml streptomycin, 0.09 M Na2SO4, 0.03 M K2SO4, and 0.014M MgCl2). The tissue was transferred to sterile conditions and enzymatically digested in dissociation medium supplied with 0.16 mg/ml L-cysteine and 70U papain (Sigma) set to pH 7.35, at 37°C for 30 min with repeated shaking. The enzyme was then inhibited with dissociation medium containing 0.1 mg/ml ovomucoid (Sigma) and 0.1 mg/ml bovine serum albumin (BSA) set to pH 7.35, at room temperature. Tissue was transferred to iced OptiMem (Life Technologies) supplied with 20 mM glucose, 0.4 mM kynurenic acid and 0.025 mM APV and mechanically dissociated until a single cell suspension was obtained. Cells were concentrated by centrifugation with 850 rpm for 5 min and filtered through a cell strainer (BD Falcon). The genetically labelled live cells were separated based on green or red fluorescence intensity using fluorescence-activated cell sorting (FACS Aria III, BD). FACS-purified cells were collected directly in lysis buffer of the RNeasy^®^ Micro Kit (Qiagen, #74004) that was used to recover total RNA according to manufacturer’s instructions. RNA quality for all samples was measured on an Agilent Bioanalyzer 2100 system. All samples with a RNA Integrity Number (RIN) > 7 were used as input to library construction.

### Library preparation and RNA sequencing

Library construction and sequencing were performed at CNAG-CRG genomics core facility (Barcelona, Spain). Briefly, cDNA multiplex libraries were prepared using SMARTer ™ Ultra Low RNA Kit v4 (Takara, #634894) and NEBNext Ultra DNA Library Prep for Illumina according to the manufacturer’s instructions (NEB, #E7645). Libraries were sequenced together in single flow cell on an Illumina HiSeq2500 platform using v4 chemistry in 1×50bp single-end mode. A minimum of 25 million reads were generated from each library.

### Bioinformatic analysis of the RNA-seq

RNA-seq analyses were performed as previously described^36^ with minor modifications: quality control of the raw data was performed with FastQC (v.0.11.7) and sequenced dataset adapters were trimmed using Cutadapt (v2.3) and Trim Galore (v0.6.1). RNA-seq reads were mapped to the Mouse genome (GRCm.38.p6/mm10) using STAR (v2.7.0d) and SAM/BAM files were further processed using SAMtools (v1.9). Aligned reads were counted and assigned to genes using Ensembl release 95 gene annotation and FeatureCounts, Subread (v1.6.4)^37^. Normalization of read counts and differential expression analyses were performed using DESeq2 (v1.22.2), Bioconductor (v3.8) in the R statistical computing and graphics platform (v3.5.1 “Feather Spray”). In the analysis of cortical astrocytes and thalamic astrocytes and neurons datasets, significantly Differentially Expressed Genes (DEGs) were identified using statistical significance threshold (Benjamini-Hochberg (BH) adjusted *P-*value < 0.1) and absolute log2 fold change (log2FC) > 0 using shrunken log2FC using the adaptive *t* prior Bayesian shrinkage estimator “apeglm”^38^. To extract enriched genes in neurons from cortex (P1) and thalamus (P0), we used previously published data from cortical neurons with GSE serie id: GSE63482^17^. Neuronal datasets from cortex were aligned from the raw sequence and counted as indicated previously. Then raw counts from cortical and thalamic neuronal transcriptomes associated to genes in the reference were normalized together using median of rations method in DeSeq2 R package. Enriched genes in cortical and thalamic neurons were obtained ranking the genes by the ratio of normalized regularized logarithm (Rlog) gene counts. Hypergeometric test (one-sided Fisher’s exact test) was performed to test independence between lists of enriched or significantly differentially expressed genes from neurons and astrocytes from different brain regions and to obtain estimated odds ratios. RNA-seq coverage tracks for selected genes were generated using IGV (v2.4.14) and plotted in a 5’ to 3’ direction. Hierarchical clustering analysis was performed using “Manhattan” distance and “Ward.2” clustering methods metrics to visualize significantly upregulated and downregulated genes. In the functional enrichment analysis of the datasets from astrocytes, a more restrictive filtering criteria was performed to detect high significantly DEG based on simultaneous threshold of BH adjusted *P*-value < 0.1 and absolute log2FC > 0.322. This analysis revealed 508 versus 444 DEGs enriched in thalamus and cortex, respectively. The Gene Ontology (GO) Over-Representation Analysis and Gene Set Enrichment Analysis (GSEA) were performed using clusterProfiler (v3.10.1)^39^. All enriched terms were considered significant at adjusted *P*-values by ‘BH’ with *P*-value cut-off < 0.01 and 0.1, in the GO Over-Representation Analysis and GSEA, respectively. The reference gene set used to perform the analysis was C5 (GO Biological Process) collection from Molecular Signatures Database (MSigDB) (v6.2).

### Bioinformatic analysis of the single-cell RNA-seq

We analysed recent work from single-cell RNA-seq (scRNA-seq) to interrogate thalamic and cortical cellular heterogeneity^18^. The sequence data is publicly available at the NCBI Sequence Read Archive (SRA) under accession SRP135960^18^. Briefly, scRNA-seq datasets (count matrices) for thalamus and cortex were downloaded from the associated wiki and processed with Seurat R package (v3.1.4)^40^. First, we performed quality control analysis that confirmed that the data were of high quality. All cells had more than 600 detected molecules (UMIs) and low percentage of reads mapped to mitochondrial transcripts. Next, data were pre-processed (log-normalization and scaling) before performing linear dimensional reduction (PCA). Graph-based clustering approach using top 30 principal components was used to identify cells populations (resolution was fixed to 0.8). FindAllMarkers function was used to identify gene markers for each cluster and to assign cell types identity to clusters. Cortical and thalamic scRNA-seq datasets were subsequently integrated as previously described^41^. The UMAP (Uniform Manifold Approximation and Projection) algorithm was used to nonlinear dimensionality reduction, visualization and exploratory analysis of the datasets. Differential expression analyses between thalamic and cortical neurons and astrocytes were performed using the *FindMarkers* function based on the non-parametric Wilcoxon rank-sum test with the following parameters (logFC.threshold = 0.1; min.pct = 0). Genes with BH adjusted *P-*value < 0.1 were considered significantly differentially expressed.

### *In utero* electroporation of StarTrack vectors

For *in utero* electroporation, a procedure previously described was followed^42^. Pregnant females (E11.5) were deeply anesthetized with isoflurane to perform laparotomies. The embryos were exposed, and the third ventricles of the embryonic brains were visualized through the uterus with an optic fiber light source. The combination of the plasmids of the StarTrack method at a final concentration of 2μg/μl were mixed with 0.1% Fast Green (Sigma) as previously described^11,20^. The plasmids used consisted on the coding sequence of 6 fluorescent proteins (EGFP, mCherry, mKusabian Orange, mTSapphire, mCerulean and EYFP) subcloned under the regulation of the GFAP or UbC promoter for targeting specifically the astrocytes or all the cell types. Each reporter gene could be directed to the cytoplasm (PB-GFAP/UbC-XFP) or to the nucleus of the cell by fusing it with the H2B histone protein (PB-GFAP/UbC-H2B-XFP). Constructs were flanked by PiggyBac sequences to be inserted into the genome of targeted cell by a PiggyBac transposase. The plasmids were injected into the third cerebral ventricle by an injector (Nanoliter 2010, WPI). For electroporation, 5 square electric pulses of 45 V and 50ms were delivered through the uterus at 950ms intervals using a square pulse electroporator (CUY21 Edit: NepaGene Co., Japan). The surgical incision was then closed, and embryos were allowed to develop until P8. In the electroporated animals with the UbC-StarTrack combination, tamoxifen was administered at P1 as previously described^20^ for removing non-integrated copies of the electroporated plasmids through the Cre recombinase system.

### Measurement of thalamic astrocytic clones

Images were acquired with an Olympus FV1000 confocal IX81 microscope/FV10-ASW software following previously defined settings^11^. All the pictures were acquired with an 20x oil immersion objective and analyzed with ImageJ software. Only electroporated animals with labelled cells in the three first order thalamic nuclei (dLG, VPM, MGv) were used. Then, only clones with at least 3 cells and with the presence of more than 1 reporter were analyzed.

### Virus production

For the production of the retrovirus, Lenti-X 293T cells (Cat. No. 632180, Clontech) were plated on 5-10 cm dishes. Encapsulation plasmids containing gag-pol and vsv-g sequences (provided by Dr. Victor Borrell) were co-transfected with the plasmid of interest using LipoD293™ (Cat. No. SL100668, Signagen). The medium was changed after 5 hours, and the virus collected after 72h using Lenti-X-concentrator (Cat. No. 631231, Clontech).

### *In vivo* viral injections

Pups at postnatal day 3 (P3) were anesthetized on ice and placed in a digital stereotaxic. The virus was injected using with an injector (Nanoliter 2010, WPI) in the thalamus or cortex through a small skull incision.

### Astrocytes primary cultures

Postnatal astroglia was cultured as previously described^43^. Briefly, after removal of the meninges, the cortices (somatosensory and visual) and the thalamus, from P4-P6 mice were dissected and dissociated mechanically in cold KREBS 1x. Subsequently, cells were centrifuged for 10 min at 1,000 rpm, re-suspended, and plated in a medium consisting of DMEM/F12 (Gibco), 3.5 mM glucose (Sigma), 10% fetal calf serum (Gibco), 5% horse serum (Gibco), 1x GlutaMax (Fisher), 100U/μl Antibiotic/Antimycotic (Fisher) and supplemented with B27 2% (Gibco), 10 ng/mL epidermal growth factor (EGF, Roche), and 10 ng/mL fibroblast growth factor 2 (FGF2, Roche). Oligodendrocyte precursor cells were removed by brusquely shaking the culture flasks several times when changing the medium after 2 or 3 days. Cells were passaged after 1 week using trypsin/EDTA (Gibco) and plated on poly-D-lysine (Sigma) glass coated coverslips at a density of 50,000-70,000 cells per coverslip (in 24-well plates; BD Biosciences) in the same medium. The vast majority of the cells (>90%) were positive for glial fibrillary acidic protein (Gfap). Nuclei-specific thalamic astrocytic cultures were performed similarly but with a few modifications. Brains were dissected out and cut in a vibratome in 300μm slices in cold KREBS to dissect the three principal sensory thalamic nuclei; dorsal-lateral geniculate (dLG) nucleus, the somatosensory ventral-posterior medial (VPM) nucleus, and the auditory ventral-medial geniculate (MGv). Thalamic nuclei were then mechanically dissociated and plated on 6-well plates and passed when confluent. Astrocytes were infected with CAG-*GFP*-IRES-*GFP*, CAG*-(Flag)Neurog2*-IRES-*DsRed*, CAG*-(Flag)Neurog2*-IRES-*TauGFP or* CAG*-Gbx2-*IRES*-DsRed* retroviruses. After 24h, the medium was changed by a differentiation medium containing DMEM/F12 (Gibco), 3.5 mM glucose (Sigma), 1x GlutaMax (Fisher), 100U/μl Antibiotic/Antimycotic (Fisher) and supplemented with B27 2% (Gibco). BDNF (Sigma) was added at 20ng/ml every fourth day during the differentiation process.

### Histology

For immunofluorescence of reprogrammed neurons in vitro, cultures were fixed with 4% paraformaldehyde (PFA) in PBS (0.01M) for 10-15 mins at room temperature. Cultures were first incubated for 1h at room temperature in a blocking solution containing 2% BSA (Sigma) and 0.15% Triton X-100 (Sigma) in PBS 0.01M. Subsequently, the cells were incubated overnight at 4°C with the primary antibodies listed in Extended Data **Table 1**. The cells were then rinsed in PBS 0.01M and incubated for 2h at room temperature with adequate secondary antibodies listed in Extended Data **Table 2**. Counterstained was performed by the fluorescent nuclear dye Dapi (Sigma-Aldrich).

For histology in postnatal brains, mice were perfused transcardially first with PBS 0.01M and PFA 4%. Brains were kept on PFA 4% overnight, embedded with agarose 3% in PBS 0.01M and cut in slices of 80μm of thickness in a vibratome (Leica). Slices were incubated for 1h at room temperature in a blocking solution containing 1% BSA and 0.3% Triton X-100 in PBS 0.01M, and subsequently, incubated overnight at 4°C with primary antibodies. For Tbr1, Ctip2, Ror*α* and Lef1 antibodies, an antigen retrieval step with sodium citrate was performed. Slices were incubated for 2 hours at room temperature with the appropriates secondary antibodies, washed, incubated with Dapi and mounted. Images were acquired with a Leica DFC550 camera into a Leica DM5000B microscope, with an Olympus FV1000 confocal IX81 microscope/FV10-ASW software or with a Zeiss confocal LSM880.

### Purification of total RNA and quantitative real-time PCR

For specific isolation of reprogrammed astrocytes, a previously published method was followed^17^ but with some modifications for cultured cells. Astrocytes from thalamus, cortex, dLG, VPM and MGv were cultured, infected with *Neurog2* retrovirus and after 10 days *in vitro* they were collected by applying Trypsin/EDTA (Gibco) to the plate, resuspended with culture medium and centrifuged. Reprogrammed astrocytes were fixed with PFA 1% for 10 minutes at 4°C, after which the PFA was quenched by adding 55μl of Glycine 1.25M per 500μl of PFA solution. Immunocytochemistry against Tuj1 and RFP was performed, and cells were separated (Tuj1+/RFP+ versus Tuj1-/RFP+) by a flow cytometer (BD FACS Aria) based on their fluorescence (see schema on Extended Data **Fig. 8**). Once the cells were collected, they were centrifuged, incubated for 3h at 50°C with lysis buffer, purified their RNA using Trizol (Fisher) and resuspended in RNase-free water.

cDNA was obtained from 1µg of total RNA using the specific protocol for First-Strand cDNA synthesis in two-step RT-PCR using the High-Capacity cDNA Reverse Transcription Kit (Fisher) and stored at −20°C. Quantitative PCR was performed in a StepOnePlus™ Real-Time PCR System (Applied Biosystems, Foster City, CA, USA) using the MicroAmp^®^ fast 96-well reaction plate (Applied Biosystems) and the Power SYBR^®^ Green PCR Master Mix (Applied Biosystems). The primers used for detecting the expression of the different genes are listed in Extended Data **Table 3**. A master mix was prepared for each primer set containing the appropriate volume of SYBR^®^ Green, primers and template cDNA. All reactions were performed in triplicate. The amplification efficiency for each primer pair and the cycle threshold (Ct) were determined automatically by the StepOne™ Software, v2.2.2 (Applied Biosystems). Transcript levels were represented relative to the *Gapdh* signal adjusting for the variability in cDNA library preparation.

### Patch-clamp recordings of iNs

For the electrophysiological analysis, astrocytes were infected with a retrovirus encoding CAG-*Neurog2*-ires-*TauGFP*. After 3 weeks, cultures were transferred to the recording chamber and were perfused with standard artificial cerebrospinal fluid (aCSF) containing (in mM): 119 NaCl, 5 KCl, 1.3 Mg SO4, 2.4 CaCl2, 1 NaH2PO4, 26 Na2HCO3 and 11 glucose. The aCSF was perfused at a rate of 2.7 ml min^-1^, continuously bubbled with a gas mixture of 95% O2+ 5% CO2 and warmed at 30-32 °C.

Somatic whole-cell recordings were made under visual control using an upright microscope (Leica DM-LFSA) and a water immersion (20 or 40x) objective. The intracellular solution contained (in mM): 130 K-gluconate, 5 KCl, 5 NaCl, 0.2 EGTA, 10 HEPES, 4 Mg-ATP and 0.4 Na-GTP, pH 7.2 adjusted with KOH; 285-295 mOsm. Recordings were obtained in current-clamp and/or voltage-clamp mode with a patch-clamp amplifier (Multiclamp 700 A, Molecular Devices, USA). No correction was made for the pipette junction potential. Voltage and current signals were filtered at 2-4 kHz and digitized at 20 kHz with a 16-bit resolution analog to digital converter (Digidata 1550-B, Axon Instruments). The generation and acquisition of pulses were controlled by pClamp 10.6 software (Axon Instruments). Patch pipettes were made from borosilicate glass (1.5 mm o.d., 0.86 mm i.d., with inner filament) and had a resistance of 4-7 MΩ when filled. Neurons in which series resistance was >30 MΩ were discarded. Quantification of intrinsic membrane properties and spontaneous neuronal activity was performed on Clampfit10.7 (Axon Instruments). The presence of putative spontaneous excitatory postsynaptic currents (sEPSCs) was assessed in voltage clamp recordings at −70 mV.

### *In silico Neurog2* binding sites determination

*In silico* analysis was performed to find out Neurog2 binding sites across the whole genome using FIMO Motif Scanning from MEME Suite (v5.0.2)^44^. Neurog2 Transcription Factor motif (NGN2_MOUSE.H11MO.0.C) from HOCOMOCO database (v11) and Mouse genome (GRCm38.p6 GenCode M18) were used to carry out this analysis. Neurog2 binding sites were annotated to genes using ChIPseeker (v1.18)^45^, Bioconductor (v3.8) in the R statistical computing and graphics platform (v3.5.1 “Feather Spray”). We retrieved genomic regions and selected binding sites (Promoters, 5’ UTR, 1st intron and 1st exon) whose location was ±3 Kb of GeneCode annotated TSSs of protein coding genes. These criteria retrieved 180611 putative Neurog2 binding sites belonging to 20478 protein coding genes. ChIP-seq coverage tracks for selected genes were generated using IGV (v2.4.14) and plotted in a 5’ to 3’ direction,, based on publicly available datasets from forebrain samples of H3K4me3 (ENCSR258YWW experiment) and H3K27me3 (ENCSR070MOK experiment) histone marks at P0 was extracted from The ENCODE Project (Extended Data Fig. **9**).

### Chromatin immunoprecipitation for H3K4me3 and H3K27me3

ChIP-assays were performed following a previously published protocol^46^. Cultured astrocytes from thalamus and cortex were collected after 1 week in vitro when reached confluent, centrifuged and resuspended to 500.000 cells approximately. Cells were fixed with 1% PFA in PBS during 10min at room temperature and quenched with 55μl of Glycine 1.25M per 500μl of PFA solution with orbital shaking. After that, cells were lysed in 300μl of SDS lysis buffer (0.5% SDS, 10mM EDTA, 50mM Tris-HCl) supplemented with protease inhibitor cocktail (Roche, 11836153001), sonicated for 10min in a Diagenode Bioruptor Pico, precleared with 30μl of washed Dynabeads (Invitrogen, 10003D) and diluted 5 times in ChIP IP buffer (20mM HEPES, 0.2M NaCl, 2mM EDTA, 0.1% Na-DOC, 1% Triton X-100, 5mg/ml BSA). 1% of each sample was kept as input. Samples were divided in 3 tubes and incubated overnight at 4°C in a rotating wheel with 2.5μg per tube of the anti-H3K4me3 (Sigma, 07-473), anti-H3K27me3 (Abcam, ab6002) or control IgG antibody. Next day, washed and saturated dynabeads were added and incubated with the samples 2h at 4°C. Dynabeads were washed 5 times with LiCl buffer (50mM HEPES, 1mM EDTA, 1% NP-40, 1% Na-DOC, 0.5M LiCl) and once with TE buffer (10mM Tris-HCl, 1mM EDTA). Antibody/chromatin complexes together with the inputs were eluted by adding 100μl of elution buffer (50mM NaHCO3, 1% SDS), 10μl of NaCl 5M, and 1μl of Proteinase K (Sigma, 3115836001) to each tube, and put on a thermomixer shaking at 1000rpm at least 2 hours at 60°C. Samples and inputs were decrosslinked by heating 15min at 95°C. Both samples and inputs were treated with RNase A (Roche, 10109142001) 30 min at 65°C and the DNA was purified with Phenol/Chloroform and ethanol precipitated. Primers used for detecting the immunoprecipitated genomic regions are listed in Extended Data **Table 4**.

### Primer design

For RNA expression analysis, Primer3 and Blast tools from NCBI webpage were used, using the accession numbers of the coding sequences of the genes of interest. For ChIP experiments, we used the information obtained from the *in silico Neurog2* binding sites analysis, and the open source information of the ENCODE project. For primers design, regions on the promoters of candidate genes that included a putative binding site for *Neurog2*, and that were enriched in H3K4me3 and H3K27me3 signal were selected.

### Quantification and Statistical analysis

Statistical analysis was carried out in GraphPad Prism (v.6™), Origin (v.8.0™) and R (v3.5.1 “Feather Spray”) statistical computing and graphics platform. Data are presented as mean and SEM. Statistical comparison between groups was performed using paired or unpaired two-tailed Student’s *t* test or Mann-Whitney U-Test non-parametric two-tailed test when data failed a Kolmogorov-Smirnov or a Shapiro Wilk normality tests. For multiple comparison analysis, a One-way ANOVA test with Holm-Sidak’s multiple comparisons test was used, and Krustal-Wallis test with Dunn’s multiple comparisons test was used when data failed a Kolmogorov-Smirnov or a Shapiro Wilk normality tests. Simple effect analysis was performed when interaction was significant. *P*-values < 0.05 were considered statistically significant and set as follows \**P* < 0.05; \*\**P* < 0.005 and \*\*\**P* < 0.0005. In the bioinformatical analysis, Differentially Expressed Genes (DEGs) were identified using statistical significance threshold (Benjamini-Hochberg (BH) adjusted *P-*value < 0.1) and set as follows *adj. *P* < 0.1; **adj. *P* < 0.01 and ***adj. *P* < 0.001. No statistical methods were used to predetermine the sample size, but our sample sizes are considered adequate for the experiments and consistent with the literature. The mice were not randomized. The investigators were blinded to sample identity.

For Fig. **1b**, PCA of astrocytes showing only the first two principal component, PC1 represents 59% variance, PC2 represents 15% variance (*n* = 4 Ctx, *n* = 4-5 each Th nucleus). For Fig. **1c**, DE analysis (adj. *P* < 0.1, log2FC > 0; 1675 DEG As-Th vs 1287 DEG As-Ctx) For Fig. **1d**, Gene Set Enrichment Analysis: Thalamus Development (GO:0021794) (NES = 1.666; *P* = 0.028; *adj. P* = 0.074), Diencephalon Development (GO:00221536) (NES = 1.889; *P* = 0.018; *adj. P* = 0.052), Cerebral Cortex Neuron Differentiation (GO:0021895) (NES = −2.119; *P* = 0.002; *adj. P* = 0.011), and Telencephalon Regionalization (GO:0021978) (NES = −1.879; *P* = 0.008; a*dj. P* = 0.029) produced with a more restrictive DE analysis (adj.*P* < 0.1, log2FC > 0.322, As-Th: 508 DEGs, As-Ctx: 444 DEGs). For Fig. **1e**, Hypergeometric test (one-sided Fisher’s exact test). As-Th enriched in Ns-Th \*\*\**P =* 3.649224e-37, OD = 4.7272; As-Th enriched in Ns-Cx ns *P =* 0.9981097, OD = 0.5754116; As-Cx enriched in Ns-Cx \*\*\**P* = 3.304775e-11, OD = 2.669471, As-Cx enriched in Ns-Th ns *P =* 0.9957097, OD = 0.5668416. Quantification was recovered from data of RNA-seq analysis of astrocytic DEGs (adj. *P* < 0.1, log2FC > 0; 1675 DEG As-Th vs 1287 DEG As-Ctx) and Top 400 neuronal genes. For Fig. **1h**, Hypergeometric test (one-sided Fisher’s exact test) from intersect between the populations of genes for the comparison of significative overexpression in As-dLG and enriched in Ns-dLG \*\*\**P =* 1.750051e-11, OD = 4.4.292546; significative overexpression in As-MGv and enriched in Ns-dLG ns *P =* 0.9956851, OD = 0.3838985; significative overexpression in As-VPM and enriched in Ns-dLG ns *P =* 0.9999685, OD = 0.5766173; significative overexpression in As-dLG and enriched in Ns-MGv ns *P =* 0.7531583, OD = 0.8598092; significative overexpression in As-MGv and enriched in Ns-MGv \*\*\**P =* 3.358423e-18, OD = 3.946944; significative overexpression in As-VPM and enriched in Ns-MGv ns *P =* 1, OD = 0.3334283; significative overexpression in As-dLG and enriched in Ns-VPM ns *P =* 0.8043838, OD = 0.7983417; significative overexpression in As-MGv and enriched in Ns-VPM ns *P =* 0.3478912, OD = 1.123736; significative overexpression in As-VPM and enriched in Ns-VPM \*\*\**P =* 1.256227e-25, OD = 2.495969. Quantification was recovered from data of RNA-seq analysis of astrocytic DEGs (adj. *P* < 0.1, log2FC > 0; 221 DEG As-dLG, 1771 DEG As-VPM, 278 DEG As-MGv) and neuronal DEGs (adj. *P* < 0.1, log2FC > 0; 705 DEG Ns-dLG, 961 DEG Ns-VPM, 1330 DEG Ns-MGv) between distinct sensory-modality thalamic nuclei.

For Fig. **2c**, Krustal-Wallis test, with Dunn’s multiple comparisons test. *n* = 5 electroporated mice. For dLG clones, *n* = 59 clones: \*\*\**P* < 0.0001; dLG (CI: 77.24%-87.74%) vs. VPM (CI: 6.932%-15.36%), \*\*\**P* < 0.0001; dLG vs. MGv (CI: 2.909%-9.824%), \*\*\**P* < 0.0001; VPM vs. MGv, ns *P* = 0.7795. For VPM clones, *n* = 179 clones \*\*\**P* < 0.0001; dLG (CI: 3.65%-7.266%) vs. VPM (CI: 84.69%-90.01%), \*\*\**P* < 0.0001; VPM vs. MGv (CI: 5.131%-9.401%), \*\*\**P* < 0.0001; dLG vs. MGv, ns *P* > 0.9999. For MGv clones, *n* = 82 clones \*\*\**P* < 0.0001; dLG (CI: 0.7806%-4.859%) vs. MGv (CI: 78.39%-88.03%), \*\*\**P* < 0.0001; VPM (CI: 8.903%-16.60%) vs. MGv, \*\*\**P* < 0.0001; dLG vs. VPM, \**P* = 0.0253. For Fig. **2g**, Krustal-Wallis test, with Dunn’s multiple comparisons test. *n* = 4 electroporated animals. For dLG mixed clones, *n* = 52 clones. For all cells, \*\*\**P* < 0.0001; dLG (Confidence interval (CI): 72.52%-81.64%) vs. VPM (CI: 13.12%-21.29%), \*\*\**P* < 0.0001; dLG vs. MGv (CI: 3.346%-7.865%), \*\*\**P* < 0.0001. For neurons, \*\*\**P* < 0.0001; dLG (CI: 64.72%-82.4%) vs. VPM (CI: 11.40%-27.25%), \*\*\**P* < 0.0001; dLG vs. MGv (CI: 3.305%-10.61%), \*\*\**P* < 0.0001. For no-neurons, \*\*\**P* < 0.0001; dLG (CI: 57.50%-78.95%) vs. VPM (CI: 14.45%-33.54%), \*\*\**P* < 0.0001; dLG vs. MGv (CI: 1.136%-14.42%), \*\*\**P* < 0.0001. For VPM clones, *n* = 71 clones. For all cells, \*\*\**P* < 0.0001; dLG (CI: 9.353%-16.29%) vs. VPM (CI: 74.28%-82.26%), \*\*\**P* < 0.0001; VPM vs. MGv (CI: 5.639%-11.73%), \*\*\**P* < 0.0001. For neurons, \*\*\**P* < 0.0001; dLG (CI: 17.88%-33.37%) vs. VPM (CI: 54.55%-71.69%), \*\*\**P* < 0.0001; VPM vs. MGv (CI: 4.817%-17.69%), \*\*\**P* < 0.0001. For no-neurons, \*\*\**P* < 0.0001; dLG (CI: 0.7925%-7.465%) vs. VPM (CI: 74.88%-90.40%), \*\*\**P* < 0.0001; VPM vs. MGv (CI: 5.491%-19.11%), \*\*\**P* < 0.0001. For MGv clones, *n* = 7 clones. For all cells, \*\*\**P* < 0.0001; VPM (CI: 6.127%-33.87%) vs. MGv (CI: 44.95%-87.91%), \*\**P* = 0.0099; dLG (CI: 1.42%-25.72%) vs. MGv, \*\**P* = 0.0011. For neurons, ns *P* = 0.7463; VPM (CI: −2.264%-68.93%) vs. MGv (CI: 0.3102%-85.40%), ns *P* > 0.9999; dLG (CI: −10.49%-58.11%) vs. MGv, ns *P* = 0.8395. For no-neurons, \**P* = 0.0167; VPM (CI: −7.477%-28.91%) vs. MGv (CI: 35.05%-107.8%), \**P* = 0.0209; dLG (CI: −16.73%-52.45%) vs. MGv, \**P* = 0.0354.

For Fig. **3c**, Mann-Whitney U-Test non-parametric two-tailed test. For Lef1, \*\**P* = 0.0065 *n* = 6 mice (265 iNs in Th, and 103 iNs in Ctx); for Ror*α*, \**P* = 0.0286 *n* = 4 mice (69 iNs in Th, and 176 iNs in Ctx); for Tbr1, \*\**P* = 0.0022 *n* = 5-6 mice (202 iNs in Th, and 109 iNs in Ctx); and for Ctip2, \*\**P* = 0.0022 *n* = 6 mice (202 iNs in Th, and 109 iNs in Ctx). For Fig. **3f**, ordinary one-way ANOVA and Holm-Sidak’s multiple comparisons test for Ror*α* (F = 13.00, Degrees of Freedom (DF) = 29, \*\*\**P* < 0.0001), Tbr1 (F = 23.56, DF = 31, \*\*\**P* < 0.0001), and Ctip2 (F = 30.70, DF = 28, \*\*\**P* < 0.0001), and Krustal-Wallis test with Dunn’s multiple comparisons test for vGlut2 (\*\*\**P* < 0.0001). For Ror*α*, Inf. As-Th vs. Inf. As-Cx, \*\*\**P* < 0.0001 *n* = 6 independent cultures; Inf. As-Th + As-Cx vs. Inf. As-Cx + As-Th, \*\*\**P* = 0.0008 *n* = 6; Inf. As-Th + Ns-Cx vs. Inf. As-Cx + Ns-Th, \*\**P* = 0.0022 *n* = 6. For Tbr1, Inf. As-Th vs. Inf. As-Cx, \*\*\**P* < 0.0001 *n* = 6; Inf. As-Th + As-Cx vs. Inf. As-Cx + As-Th, \*\*\**P* < 0.0001 *n* = 6; Inf. As-Th + Ns-Cx vs. Inf. As-Cx + Ns-Th, \*\*\**P* < 0.0001 *n* = 6. For Ctip2, Inf. As-Th vs. Inf. As-Cx, \*\*\**P* < 0.0001 *n* = 6; Inf. As-Th + As-Cx vs. Inf. As-Cx + As-Th, \*\*\**P* < 0.0001 *n* = 6; Inf. As-Th + Ns-Cx vs. Inf. As-Cx + Ns-Th, \*\*\**P* < 0.0001 *n* = 6. For vGlut2, Inf. As-Th vs. Inf. As-Cx, \*\*\**P* = 0.0003 *n* = 6; Inf. As-Th + As-Cx vs. Inf. As-Cx + As-Th, \**P* = 0.0437 *n* = 7; Inf. As-Th + Ns-Cx vs. Inf. As-Cx + Ns-Th, \**P* = 0.0239 *n* = 5. For Fig. **3h**, Ordinary one-way ANOVA test with Holm-Sidak’s multiple comparisons test. *Sp9*, F = 8.924, DF = 32, \*\*\**P* = 0.0008; dLG vs. VPM, \*\*\**P* = 0.0007 *n* = 12 independent cultures; dLG vs. MGv, \*\**P* = 0.0034 *n* = 12. *Hs6st2*, F = 5.128, DF = 25, \**P* = 0.0136; dLG vs. VPM, \*\**P* = 0.0093 *n* = 10; dLG vs. MGv, \**P* = 0.0331 *n* = 10. *Crabp2*, F = 4.702, DF = 24, \**P* = 0.0189; MGv vs. dLG, \**P* = 0.0341 *n* = 10; MGv vs. VPM, \**P* = 0.0147 *n* = 10. *Tshz1*, F = 10.97, DF = 37, \*\*\**P* = 0.0002; MGv vs. dLG, \*\*\**P* = 0.0006 *n* = 14; MGv vs. VPM, \*\*\**P* = 0.0003 *n* = 14. *Cck*, F = 5.409, DF = 30, \*\**P* = 0.0099; VPM vs. dLG, \*\**P* = 0.0064 *n* = 12; VPM vs. MGv, \**P* = 0.0393 *n* = 12.

For Fig. **4b**, Ordinary one-way ANOVA test with Holm-Sidak’s multiple comparisons test; for *Gbx2* in Th, F = 39.71, \*\*\**P* < 0.0001; Th basal (*n* = 19) vs. Th + Neurog2 (*n* = 14), ns *P* = 0.9579, t = 0.05318, DF = 36; Th basal vs. Th + Neurog2 + Gbx2 (*n* = 6), \*\*\**P* < 0.0001, t = 8.429, DF = 36; Th + Neurog2 vs. Th + Neurog2 + Gbx2, \*\*\**P* < 0.0001, t = 8.128, DF = 36. For Ctx,, F = 167.4, \*\*\**P* < 0.0001; Ctx basal (*n* = 14) vs. Ctx + Neurog2 (*n* = 14), ns *P* = 0.9831, t = 0.02134, DF = 31; Ctx basal vs. Ctx + Neurog2 + Gbx2 (*n* = 6), \*\*\**P* < 0.0001, t = 16.88, DF = 31; Ctx + Neurog2 vs. Ctx + Neurog2 + Gbx2, \*\*\**P* < 0.0001, t = 16.86, DF = 31. *Pou2f2*, for Th, F = 20.15, \*\*\**P* < 0.0001; Th basal (*n* = 12) vs. Th + Neurog2 (*n* = 14), \**P* = 0.0386, t = 2.163, DF = 30; Th basal vs. Th + Neurog2 + Gbx2 (*n* = 7), \*\*\**P* < 0.0001, t = 6.307, DF = 30; Th + Neurog2 vs. Th + Neurog2 + Gbx2, \*\*\**P* = 0.0001, t = 4.642, DF = 30. For Ctx, F = 11.79, \*\*\**P* = 0.0001; Ctx basal (*n* = 14) vs. Ctx + Neurog2 (*n* = 12), ns *P* = 0.6091, t = 0.5164, DF = 32; Ctx basal vs. Ctx + Neurog2 + Gbx2 (*n* = 9), \*\*\**P* = 0.0002, t = 4.589, DF = 32; Ctx + Neurog2 vs. Ctx + Neurog2 + Gbx2, \*\*\**P* = 0.0007, t = 3.986, DF = 32. *Tbr1*, for Th, F = 0.2125, ns *P* = 0.8095; Th basal (*n* = 20) vs. Th + Neurog2 (*n* = 14), ns *P* = 0.8900, t = 0.6478, DF = 39; Th basal vs. Th + Neurog2 + Gbx2 (*n* = 8), ns *P* = 0.9203, t = 0.1537, DF = 39; Th + Neurog2 vs. Th + Neurog2 + Gbx2, ns *P* = 0.9023, t = 0.3642, DF = 39. For Ctx, F = 5.79, \*\**P* = 0.0062; Ctx basal (*n* = 20) vs. Ctx + Neurog2 (*n* = 15), \*\**P* = 0.0076, t = 3.222, DF = 40; Ctx basal vs. Ctx + Neurog2 + Gbx2 (*n* = 8), ns *P* = 0.9186, t = 0.1029, DF = 40; Ctx + Neurog2 vs. Ctx + Neurog2 + Gbx2, \**P* = 0.0403, t = 2.416, DF = 40. *Ctip2*, for Th, F = 15.57, \*\*\**P* < 0.0001; Th basal (*n* = 19) vs. Th + Neurog2 (*n* = 14), \*\*\**P* < 0.0001, t = 5.452, DF = 37; Th basal vs. Th + Neurog2 + Gbx2 (*n* = 7), ns *P* = 0.4738, t = 0.7238, DF = 37; Th + Neurog2 vs. Th + Neurog2 + Gbx2, \*\**P* = 0.0028, t = 3.457, DF = 37. For Ctx, F = 4.681, \**P* = 0.0154; Ctx basal (*n* = 20) vs. Ctx + Neurog2 (*n* = 12), \**P* = 0.018, t = 2.913, DF = 37; Ctx basal vs. Ctx + Neurog2 + Gbx2 (*n* = 8), ns *P* = 0.9460, t = 0.06815, DF = 37; Ctx + Neurog2 vs. Ctx + Neurog2 + Gbx2, \**P* = 0.05, t = 2.268, DF = 37. *Rorα*, for Th, F = 0.7022, ns *P* = 0.5015; Th basal (*n* = 22) vs. Th + Neurog2 (*n* = 14), ns *P* = 0.5676, t = 1.183, DF = 40; Th basal vs. Th + Neurog2 + Gbx2 (*n* = 7), ns *P* = 0.8707, t = 0.4299, DF = 40; Th + Neurog2 vs. Th + Neurog2 + Gbx2, ns *P* = 0.8707, t = 0.4707, DF = 40. For Ctx, F = 0.05697, ns *P* = 0.9447; Ctx basal (*n* = 22) vs. Ctx + Neurog2 (*n* = 14), ns *P* = 0.9827, t = 0.1035, DF = 41; Ctx basal vs. Ctx + Neurog2 + Gbx2 (*n* = 8), ns *P* = 0.9827, t = 0.2709, DF = 41; Ctx + Neurog2 vs. Ctx + Neurog2 + Gbx2, ns *P* = 0.9827, t = 0.3321, DF = 41. *Lef1*, for Th, F = 0.2178, ns *P* = 0.8053; Th basal (*n* = 21) vs. Th + Neurog2 (*n* = 14), ns *P* = 0.8893, t = 0.6494, DF = 40; Th basal vs. Th + Neurog2 + Gbx2 (*n* = 8), ns *P* = 0.9353, t = 0.3267, DF = 40; Th + Neurog2 vs. Th + Neurog2 + Gbx2, ns *P* = 0.9353, t = 0.1993, DF = 40. For Ctx, F = 0.4896, ns *P* = 0.6164; Ctx basal (*n* = 22) vs. Ctx + Neurog2 (*n* = 14), ns *P* = 0.7079, t = 0.9725, DF = 41; Ctx basal vs. Ctx + Neurog2 + Gbx2 (*n* = 8), ns *P* = 0.8896, t = 0.1396, DF = 41; Ctx + Neurog2 vs. Ctx + Neurog2 + Gbx2, ns *P* = 0.7871, t = 0.6201, DF = 41. *Fezf2*, for Th, F = 21.11, \*\*\**P* < 0.0001; Th basal (*n* = 17) vs. Th + Neurog2 (*n* = 10), \*\*\**P* < 0.0001, t = 5.764, DF = 32; Th basal vs. Th + Neurog2 + Gbx2 (*n* = 8), \*\*\**P* < 0.0001, t = 4.800, DF = 32; Th + Neurog2 vs. Th + Neurog2 + Gbx2, ns *P* = 0.6177, t = 0.5040, DF = 32. For Ctx, F = 0.2050, ns *P* = 0.8157; Ctx basal (*n* = 18) vs. Ctx + Neurog2 (*n* = 10), ns *P* = 0.9051, t = 0.3393, DF = 33; Ctx basal vs. Ctx + Neurog2 + Gbx2 (*n* = 8), ns *P* = 0.9051, t = 0.3998, DF = 33; Ctx + Neurog2 vs. Ctx + Neurog2 + Gbx2, ns *P* = 0.8938, t = 0.6402, DF = 33. *vGlut2*; for Th, F = 4.011, \**P* = 0.0261; Th basal (*n* = 21) vs. Th + Neurog2 (*n* = 13), \**P* = 0.0228, t = 2.813, DF = 39; Th basal vs. Th + Neurog2 + Gbx2 (*n* = 8), ns *P* = 0.4043, t = 1.224, DF = 39; Th + Neurog2 vs. Th + Neurog2 + Gbx2, ns *P* = 0.4043, t = 1.077, DF = 39. For Ctx, F = 3.454, \**P* = 0.0427; Ctx basal (*n* = 20) vs. Ctx + Neurog2 (*n* = 9), ns *P* = 0.4808, t = 0.7127, DF = 35; Ctx basal vs. Ctx + Neurog2 + Gbx2 (*n* = 9), ns *P* = 0.0702, t = 2.184, DF = 35; Ctx + Neurog2 vs. Ctx + Neurog2 + Gbx2, ns *P* = 0.0550, t = 2.466, DF = 35.

In Fig. **4d**, for the log2 of the Ratio H3K4me3/HeK27me3, unpaired student’s t-test two-tailed test; *Gbx2*, Th (*n* = 14) vs. Ctx (*n* = 12) \*\*\**P* < 0.0001, t = 8.037, df = 24; *Rorα*, Th (*n* = 23) vs. Ctx (*n* = 22) \**P* = 0.0450, t = 2.065, df = 43; *Lef1*, Th (*n* = 20) vs. Ctx (*n* = 21) \**P* = 0.0126, t = 2.616, df = 39; *Fezf2*, Th (*n* = 10) vs. Ctx (*n* = 10) ns *P* = 0.3111, t = 1.042, df = 18; *vGlut2*, Th (*n* = 16) vs. Ctx (*n* = 16) ns *P* = 0.2250, t = 1.239, df = 30; *Pou2f2*, Th (*n* = 16) vs. Ctx (*n* = 18) ns *P* = 0.5076, t = 0.6700, df = 32; *Tbr1*, Th (*n* = 19) vs. Ctx (*n* = 21) \**P* = 0.0152, t = 2.542, df = 38; *Ctip2*, Th (*n* = 17) vs. Ctx (*n* = 18) \*\**P* = 0.0013, t = 3.524, df = 33. For the expression levels (1/ΔCt), unpaired student’s t-test two-tailed test; *Gbx2*, Th (*n* = 19) vs. Ctx (*n* = 14) \*\*\**P* < 0.0001, t = 9.066, df = 31; *Rorα*, Th (*n* = 22) vs. Ctx (*n* = 14) \**P* = 0.0216, t = 2.409, df = 34; *Lef1*, Th (*n* = 22) vs. Ctx (*n* = 22) \*\*\**P* < 0.0001, t = 6.388, df = 42; *Fezf2*, Th (*n* = 10) vs. Ctx (*n* = 10) \*\**P* = 0.0028, t = 3.458, df = 18; *vGlut2*, Th (*n* = 21) vs. Ctx (*n* = 14) \**P* = 0.0298, t = 2.271, df = 33; *Pou2f2*, Th (*n* = 17) vs. Ctx (*n* = 11) \**P* = 0.0118, t = 2.708, df = 26; *Tbr1*, Th (*n* = 13) vs. Ctx (*n* = 20) ns *P* = 0.3033, t = 1.047, df = 31; *Ctip2*, Th (*n* = 14) vs. Ctx (*n* = 14) ns *P* = 0.1874, t = 3.524, df = 26.

In Fig. **4e**, for the epigenetics, Ordinary one-way ANOVA test with Holm-Sidak’s multiple comparisons test. For *Sp9*, F = 6.486, \*\**P* = 0.0036; dLG vs. VPM, \*\**P* = 0.0018, *n* = 14, t = 3.587, DF = 41; dLG vs. MGv, \**P* = 0.0394 *n* = 14-16, t = 2.128, DF = 41. *Hs6st2*, F = 4.188, \**P* = 0.0215; dLG vs. VPM, \**P* = 0.0164, *n* = 13-18, t = 2.764, DF = 45; dLG vs. MGv, \**P* = 0.0268, *n* = 13-17, t = 2.289, DF = 45. *Crabp2*, F = 4.794, \**P* = 0.0132; MGv vs. dLG, \**P* = 0.0409, *n* = 18-12, t = 2.108, DF = 43; MGv vs. VPM, \**P* = 0.01, *n* = 16-18, t = 2.958, DF = 43. *Tshz1*, F = 5.125, \**P* = 0.0106; MGv vs. dLG, \**P* = 0.0355, *n* = 13-15, t = 2.178, DF = 39; MGv vs. VPM, \*\**P* = 0.0072, *n* = 14-15, t = 3.098, DF = 39. *Cck*, F = 5.489, \*\**P* = 0.0076; VPM vs. dLG, \**P* = 0.0227, *n* = 13-16, t = 2.365, DF = 42; VPM vs. MGv, \*\**P* = 0.0058, *n* = 16, t = 3.164, DF = 42. For *in vivo* basal expression, data from the RNA-seq analysis of the astrocytes (adjusted *P* < 0.1, log2FC > 0.322) from the three thalamic nuclei was used.

For Extended Data Fig. **1b**, mean levels of expression are shown with black line over light green region, while standard error mean is represented with the dark green area. In Extended Data Fig. **1c**, DE analysis. (adj. *P* < 0.1, log2FC > 0; 1675 DEG As-Th vs 1287 DEG As-Ctx). For Extended Data Fig. **1e** and **1f** was used a more restrictive DE analysis (adj.*P* < 0.1, log2FC > 0.322, As-Th: 508 DEGs, As-Ctx: 444 DEGs).

For Extended Data Fig. **2e**, Pearson Correlation Coefficient, *R* = 0.9750593 (*T* = 121.83, df = 769, \*\*\**P <* 2.2e-16).

For Extended Data Fig. **4c** and **4d**, for the differential expression of astrocytes, Wilcoxon rank-sum test (adj.*P* < 0.1, log2FC > 0.1, As-Th: 549 DEGs, As-Ctx: 1106 DEGs). For the differential expression of neurons, Wilcoxon rank-sum test (adj.*P* < 0.1, log2FC > 0.1, Ns-Th: 2425 DEGs, Ns-Ctx: 1845 DEGs).

For Extended Data Fig. **5a**, PCA showing only the first two principal component in astrocytes of the three thalamic nuclei, PC1 represents 41% variance, PC2 represents 24% variance (*n* = 4-5 replicates each). In Extended Data **5b**, DE analysis DEGs (adj. P < 0.1, log2FC > 0; 221 DEG As-dLG, 1771 DEG As-VPM, 278 DEG As-MGv). For Extended Data Fig. **5c**, PCA showing only the first two principal component, PC1 represents 47% variance, PC2 represents 32% variance (*n* = 3-4 replicates each). In Extended Data Fig. **5d**, DE analysis (adj. *P* < 0.1, log2FC > 0; 705 DEGs Ns-dLG, 961 DEGs Ns-VPM, 1300 DEGs Ns-MGv).

For Extended Data Fig. **6b**, left graph, ordinary one-way ANOVA and Holm-Sidak’s multiple comparisons test, F = 0.07668, ns *P* = 0.9266; dLG vs. VPM ns *P* = 0.9737, dLG vs. MGv ns *P* = 0.9737, MGv vs. VPM ns *P* = 0.9737, *n* = 5; right graph, ordinary one-way ANOVA and Holm-Sidak’s multiple comparisons test, F = 0.3985, ns *P* = 0.6799; dLG vs. VPM ns *P* = 0.8286, dLG vs. MGv ns *P* = 0.7913, MGv vs. VPM ns *P* = 0.7913, *n* = 5 electroporated animals. In Extended Data Fig. **6c**, left graph, *n* = 59 dLG clones, *n* = 179 VPM clones, *n* = 82 MGv clones. In right graph, Krustal-Wallis test with Dunn’s multiple comparisons test. For dLG clones, *P* > 0.9999 in dLG; *P* > 0.9999 in VPM, and *P* = 0.6773 in MGv, *n* = 43 clones with 3-10 cells and *n* = 16 clones with >10 cells. For VPM clones, *P* = 0.6386 in dLG; *P* > 0.9999 in VPM, and *P* = 0.0976 in MGv, *n* = 137 clones with 3-10 cells and *n* = 44 clones with >10 cells. For MGv clones, *P* = 0.4436 in dLG; *P* > 0.9999 in VPM, and *P* > 0.9999 in MGv, *n* = 66 clones with 3-10 cells and *n* = 15 clones with >10 cells. In Extended Data Fig. **6e**, Mann-Whitney U-Test non-parametric two-tailed test (*n* = 128 clones); Neurons vs. Non-neurons, ns *P* = 0.3112. In Extended Data Fig. **6f**, Krustal-Wallis test, with Dunn’s multiple comparisons test. *n* = 4 electroporated animals. For dLG neuronal clones, *n* = 61 clones, \*\*\**P* < 0.0001; dLG (Confidence interval (CI): 80.53%-88.87%) vs. VPM (CI: 5.962%-13.56%), \*\*\**P* < 0.0001; dLG vs. MGv (CI: 3.069%-8.002%), \*\*\**P* < 0.0001. For dLG non-neuronal clones, *n* = 14 clones, \*\*\**P* < 0.0001; dLG (CI: 100%-100%) vs. VPM (CI: 0%-0%), \*\*\**P* < 0.0001; dLG vs. MGv (CI: 0%-0%), \*\*\**P* < 0.0001. For VPM neuronal clones, *n* = 7 clones, \*\*\**P* = 0.0007; dLG (CI: 0.856%-37.24%) vs. VPM (CI: 45.53%-82.08%), \*\**P* = 0.0081; dLG vs. MGv (CI: −0.13%-34.42%), \*\*\**P* = 0.0039. For VPM non-neuronal clones, *n* = 25 clones, \*\*\**P* < 0.0001; dLG (CI: −0.7909%-7.458%) vs. VPM (CI: 92.54%-100.8%), \*\*\**P* < 0.0081; dLG vs. MGv (CI: 0%-0%), \*\*\**P* < 0.0001. For MGv neuronal clones, *n* = 4 clones, \*\**P* = 0.0052; VPM (CI: −18.95%-60.61%) vs. MGv (CI: 39.39%-118.9%), ns *P* = 0.0917; dLG (CI: 0%-0%) vs. MGv, \*\**P* = 0.0077. For MGv non-neuronal clones, *n* = 3 clones, \**P* = 0.0357; VPM (CI: 0%-0%) vs. MGv (CI: 100%-100%), \**P* = 0.0286; dLG (CI: 0%-0%) vs. MGv, \*\**P* = 0.0286.

For Extended Data Fig. **7b**, in Th, \*\**P* = 0.0079 *n* = 5 injected mice (427 cells with control and 572 cells with *Neurog2* virus); and in Cx, \*\**P* = 0.0043, *n* = 5 injected mice (362 cells with control and 292 cells with *Neurog2* virus).

For Extended Data Fig. **8d**, for *Gfap*, two-tailed unpaired Student’s t-test, \*\**P* = 0.0014 *n* = 8; for *Neurog2*, ordinary one-way ANOVA test F = 23.41, \*\*\**P* < 0.0001; Holm-Sidak’s multiple comparisons test, Tuj1+/RFP+ vs. RFP+, \*\*\**P* = 0.0006; Tuj1+/RFP+ vs. -/-, \*\*\**P* < 0.0001; RFP+ vs. -/-, \**P* = 0.0116; *n* = 12 cultures. For Extended Data Fig. **8e**, Student’s t-test or Mann-Whitney U-Test non-parametric two-tailed test; *vGlut2*, \*\**P* = 0.0058 *n* = 16; *Rorα*, \*\**P* = 0.0027 *n* = 17; *Gbx2*, \*\**P* = 0.0065, *n* = 7; *Pou2f2*, \**P* = 0.0181, *n* = 5; *Lef1*, \*\**P* = 0.0016, *n* = 7; *Tbr1*, \*\**P* = 0.0078, *n* = 11; *Ctip2*, \*\**P* = 0.0079, *n* = 5.

For Extended Data Fig. **10b**, for *Gapdh* and *Cdx2* upper graphs, two-tailed unpaired Student’s t-test; for *Gapdh* \*\**P* = 0.0065 *n* = 9; and for *Cdx2*, ns *P* = 0.3406 *n* = 9. In lower graph, *Gapdh* vs. *Cdx2*, Mann-Whitney U-Test non-parametric two-tailed test, \*\*\**P* < 0.0001 *n* = 8. For *Gbx2*, two-tailed paired Student’s t-test: Th \*\*\**P* < 0.0001 *n* = 14, Ctx \*\**P* = 0.0049 *n* = 12; for *Rorα*, two-tailed Wilcoxon matched-pairs test: Th \*\*\**P* < 0.0001 *n* = 23, Ctx \**P* = 0.019 *n* = 22; for *Lef1*, two-tailed paired Student’s t-test: Th ns *P* = 0.0563 *n* = 18, Ctx ns *P* = 0.211 *n* = 23; for *Fezf2*, two-tailed paired Student’s t-test: Th ns *P* = 0.2506 *n* = 9, Ctx ns *P* = 0.3506 *n* = 10; for *vGlut2*, two-tailed Wilcoxon matched-pairs test: Th \**P* = 0.0214 *n* = 16, Ctx \*\*\**P* = 0.0003 *n* = 16; for *Pou2f2*, two-tailed Wilcoxon matched-pairs test: Th ns *P* = 0.7819 *n* = 17, Ctx ns *P* = 0.2121 *n* = 18; for *Tbr1*, two-tailed paired Student’s t-test: Th \*\*\**P* = 0.0003 *n* = 19, Ctx ns *P* = 0.1748 *n* = 21; for *Ctip2*, two-tailed paired Student’s t-test: Th \**P* = 0.0156 *n* = 17, Ctx ns *P* = 0.1471 *n* = 18. For Extended Data Fig. **10d**, *Sp9*, two-tailed paired Student’s t-test: dLG, \*\**P* = 0.0045 *n* = 14, VPM \*\*\**P* = 0.0001 *n* = 14 and MGv \*\*\**P* < 0.0001 *n* = 16; *Hs6st2*, two-tailed paired Student’s t-test: dLG, \*\**P* = 0.0031 *n* = 14, VPM \*\*\**P* = 0.0001 *n* = 18 and MGv \*\*\**P* < 0.0001 *n* = 18; *Cck*, two-tailed paired Student’s t-test: dLG, \*\*\**P* < 0.0001 *n* = 15, VPM \*\*\**P* < 0.0001 *n* = 18 and MGv \*\*\**P* < 0.0001 *n* = 18; *Crabp2*, two-tailed Wilcoxon matched-pairs test: dLG, \**P* = 0.0245 *n* = 14, VPM ns *P* = 0.0987 *n* = 18 and MGv \*\*\**P* < 0.0001 *n* = 19; *Tshz1*, two-tailed Wilcoxon matched-pairs test: dLG, \*\*\**P* < 0.0001 *n* = 13, VPM \*\*\**P* = 0.0001 *n* = 14 and MGv \*\*\**P* < 0.0001 *n* = 16.

## References

1. Metzis, V. et al. Nervous System Regionalization Entails Axial Allocation before Neural Differentiation. Cell 175, 1105–1118.e17 (2018).

2. Mi, D. et al. Early emergence of cortical interneuron diversity in the mouse embryo. Science 360, 81–85 (2018).

3. Mayer, C. et al. Developmental diversification of cortical inhibitory interneurons. Nature 555, 457–462 (2018).

4. Telley, L. et al. Temporal patterning of apical progenitors and their daughter neurons in the developing neocortex. Science 364, eaav2522 (2019).

5. Lodato, S. & Arlotta, P. Generating neuronal diversity in the mammalian cerebral cortex. Annu. Rev. Cell Dev. Biol. 31, 699–720 (2015).

6. Jessell, T. M. Neuronal specification in the spinal cord: inductive signals and transcriptional codes. Nat. Rev. Genet. 1, 20–29 (2000).

7. Greig, L. C., Woodworth, M. B., Galazo, M. J., Padmanabhan, H. & Macklis, J. D. Molecular logic of neocortical projection neuron specification, development and diversity. Nat. Rev. Neurosci. 14, 755–769 (2013).

8. Nowakowski, T. J. et al. Spatiotemporal gene expression trajectories reveal developmental hierarchies of the human cortex. Science 358, 1318–1323 (2017).

9. Rowitch, D. H. & Kriegstein, A. R. Developmental genetics of vertebrate glial-cell specification. Nature 468, 214–222 (2010).

10. Hochstim, C., Deneen, B., Lukaszewicz, A., Zhou, Q. & Anderson, D. J. Identification of positionally distinct astrocyte subtypes whose identities are specified by a homeodomain code. Cell 133, 510–522 (2008).

11. García-Marqués, J. & López-Mascaraque, L. Clonal identity determines astrocyte cortical heterogeneity. Cereb. Cortex 23, 1463–1472 (2013).

12. Bayraktar, O. A. et al. Astrocyte layers in the mammalian cerebral cortex revealed by a single-cell in situ transcriptomic map. Nat. Neurosci. 8, 427 (2020).

13. Batiuk, M. Y. et al. Identification of region-specific astrocyte subtypes at single cell resolution. Nat Commun 11, 1220–15 (2020).

14. Khakh, B. S. & Deneen, B. The Emerging Nature of Astrocyte Diversity. Annu. Rev. Neurosci. 42, 187–207 (2019).

15. Ge, W.-P., Miyawaki, A., Gage, F. H., Jan, Y. N. & Jan, L. Y. Local generation of glia is a major astrocyte source in postnatal cortex. Nature 484, 376–380 (2012).

16. Gezelius, H. et al. Genetic Labeling of Nuclei-Specific Thalamocortical Neurons Reveals Putative Sensory-Modality Specific Genes. Cereb. Cortex 27, 5054–5069 (2017).

17. Molyneaux, B. J. et al. DeCoN: genome-wide analysis of in vivo transcriptional dynamics during pyramidal neuron fate selection in neocortex. Neuron 85, 275–288 (2015).

18. Zeisel, A. et al. Molecular Architecture of the Mouse Nervous System. Cell 174, 999–1014.e22 (2018).

19. Shi, W. et al. Ontogenetic establishment of order-specific nuclear organization in the mammalian thalamus. Nat. Neurosci. 20, 516–528 (2017).

20. Figueres-Oñate, M., García-Marqués, J. & López-Mascaraque, L. UbC-StarTrack, a clonal method to target the entire progeny of individual progenitors. Sci Rep 6, 33896–13 (2016).

21. Gascón, S. et al. Identification and Successful Negotiation of a Metabolic Checkpoint in Direct Neuronal Reprogramming. Cell Stem Cell 18, 396–409 (2016).

22. Mallika, C., Guo, Q. & Li, J. Y. H. Gbx2 is essential for maintaining thalamic neuron identity and repressing habenular characters in the developing thalamus. Dev. Biol. 407, 26–39 (2015).

23. Bluske, K. K. et al. β-Catenin signaling specifies progenitor cell identity in parallel with Shh signaling in the developing mammalian thalamus. Development 139, 2692–2702 (2012).

24. Lodato, S. et al. Gene co-regulation by Fezf2 selects neurotransmitter identity and connectivity of corticospinal neurons. Nat. Neurosci. 17, 1046–1054 (2014).

25. Mihalas, A. B. & Hevner, R. F. Control of Neuronal Development by T-Box Genes in the Brain. Curr. Top. Dev. Biol. 122, 279–312 (2017).

26. Guo, Q. & Li, J. Y. H. Defining developmental diversification of diencephalon neurons through single cell gene expression profiling. Development 146, dev174284 (2019).

27. Hirabayashi, Y. et al. Polycomb limits the neurogenic competence of neural precursor cells to promote astrogenic fate transition. Neuron 63, 600–613 (2009).

28. Hirabayashi, Y. & Gotoh, Y. Epigenetic control of neural precursor cell fate during development. Nat. Rev. Neurosci. 11, 377–388 (2010).

29. Magnusson, J. P. et al. A latent neurogenic program in astrocytes regulated by Notch signaling in the mouse. Science 346, 237–241 (2014).

30. Nato, G. et al. Striatal astrocytes produce neuroblasts in an excitotoxic model of Huntington’s disease. Development 142, 840–845 (2015).

31. Mattugini, N. et al. Inducing Different Neuronal Subtypes from Astrocytes in the Injured Mouse Cerebral Cortex. Neuron 103, 1086–1095.e5 (2019).

32. Qian, H. et al. Therapeutic Reversal of Chemically Induced Parkinson Disease by Converting Astrocytes into Nigral Neurons. bioRxiv 2020.04.06.028084 (2020).

33. Tamamaki, N. et al. Green fluorescent protein expression and colocalization with calretinin, parvalbumin, and somatostatin in the GAD67-GFP knock-in mouse. J. Comp. Neurol. 467, 60–79 (2003).

34. Chen, L., Guo, Q. & Li, J. Y. H. Transcription factor Gbx2 acts cell-nonautonomously to regulate the formation of lineage-restriction boundaries of the thalamus. Development 136, 1317–1326 (2009).

35. Antón-Bolaños, N. et al. Prenatal activity from thalamic neurons governs the emergence of functional cortical maps in mice. Science 364, 987–990 (2019).

36. Scandaglia, M. et al. Loss of Kdm5c Causes Spurious Transcription and Prevents the Fine-Tuning of Activity-Regulated Enhancers in Neurons. Cell Rep 21, 47–59 (2017).

37. Liao, Y., Smyth, G. K. & Shi, W. featureCounts: an efficient general purpose program for assigning sequence reads to genomic features. Bioinformatics 30, 923–930 (2014).

38. Zhu, A., Ibrahim, J. G. & Love, M. I. Heavy-tailed prior distributions for sequence count data: removing the noise and preserving large differences. Bioinformatics 35, 2084–2092 (2019).

39. Yu, G., Wang, L.-G., Han, Y. & He, Q.-Y. clusterProfiler: an R package for comparing biological themes among gene clusters. OMICS 16, 284–287 (2012).

40. Stuart, T. et al. Comprehensive Integration of Single-Cell Data. Cell 177, 1888–1902.e21 (2019).

41. Butler, A., Hoffman, P., Smibert, P., Papalexi, E. & Satija, R. Integrating single-cell transcriptomic data across different conditions, technologies, and species. Nat. Biotechnol. 36, 411–420 (2018).

42. Moreno-Juan, V. et al. Prenatal thalamic waves regulate cortical area size prior to sensory processing. Nat Commun 8, 14172–14 (2017).

43. Heinrich, C. et al. Generation of subtype-specific neurons from postnatal astroglia of the mouse cerebral cortex. Nat Protoc 6, 214–228 (2011).

44. Bailey, T. L. et al. MEME SUITE: tools for motif discovery and searching. Nucleic Acids Res. 37, W202–8 (2009).

45. Yu, G., Wang, L.-G. & He, Q.-Y. ChIPseeker: an R/Bioconductor package for ChIP peak annotation, comparison and visualization. Bioinformatics 31, 2382–2383 (2015).

46. Gillotin, S. & Guillemot, F. Micro-chromatin Immunoprecipation (μChIP) Protocol for Real-time PCR Analysis of a Limited Amount of Cells. Bio Protoc 6, (2016).

