## Extended Data for "Astrocytes and neurons share brain region-specific transcriptional signatures"

**Supplementary Materials**

Extended Data Figures 1 to 10

Extended Data Tables 1 to 4

### Extended Data Figures

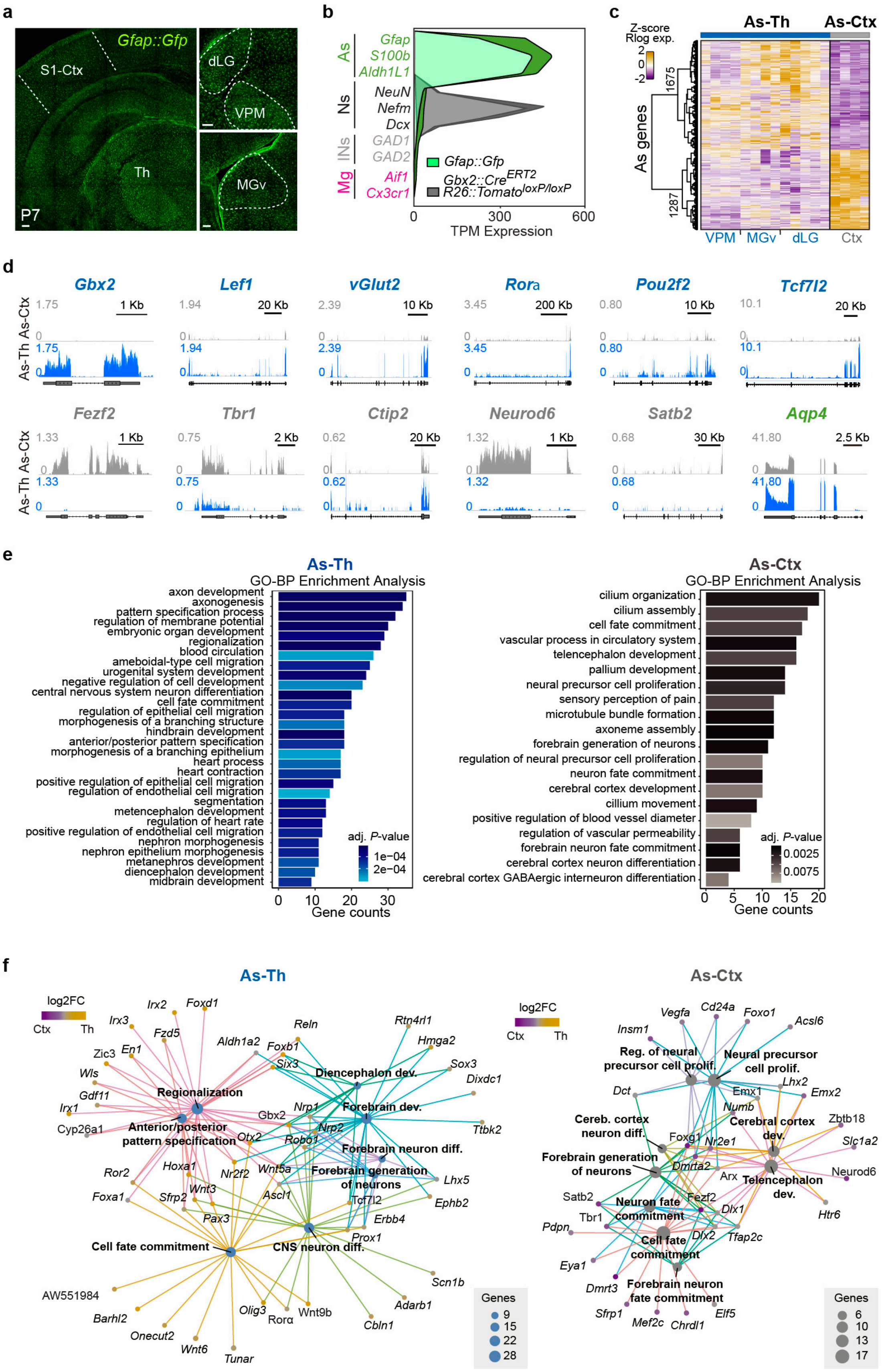

**Extended Data Fig. 1 | Differential gene expression in thalamic versus cortical astrocytes.** **a,** Images showing the thalamus and cortex of a *Gfap::Gfp* mouse at P7. **b,** Expression levels (Transcripts Per Million, TPM) of known astrocytes (As), neuronal (Ns), interneurons (INs) and microglia (Mg) genes in the purified astrocytes from a *Gfap::Gfp* mouse and purified neurons from a *Gbx2-Cre::Tomato-floxed* mouse. **c,** Heatmap of Z-score normalized regularized logarithm (Rlog) expression and unbiased clustering of significantly differentially expressed genes (DEGs) between thalamic (As-Th) and cortical astrocytes (As-Ctx). Each row represents a gene, the columns are biological replicates and the color-code represents the normalized expression for upregulated genes in yellow versus downregulated genes in purple. As-Th include dLG ($n$ = 5 samples), VPM ($n$ = 4) and MGv ($n$ = 4) and As-Ctx ($n$ = 4). **d,** Genomic tracks display normalized coverage RNA-seq data for specific significantly DEGs between thalamic and cortical astrocytes. **e**, Gene ontology (GO) biological process (BP) enrichment analysis of significantly DEGs in thalamic and cortical astrocytes. **f,** Gene ontology networks. The size of every node (enriched term) represents the number of genes enriched and the color-code (yellow, high expression; purple, low expression) corresponds to the log2FC in DE analysis. In **a,** Scale bars, 100μm. Data in **b** are mean levels of expression shown with black line over light green region, while standard error mean is represented with the dark green area.

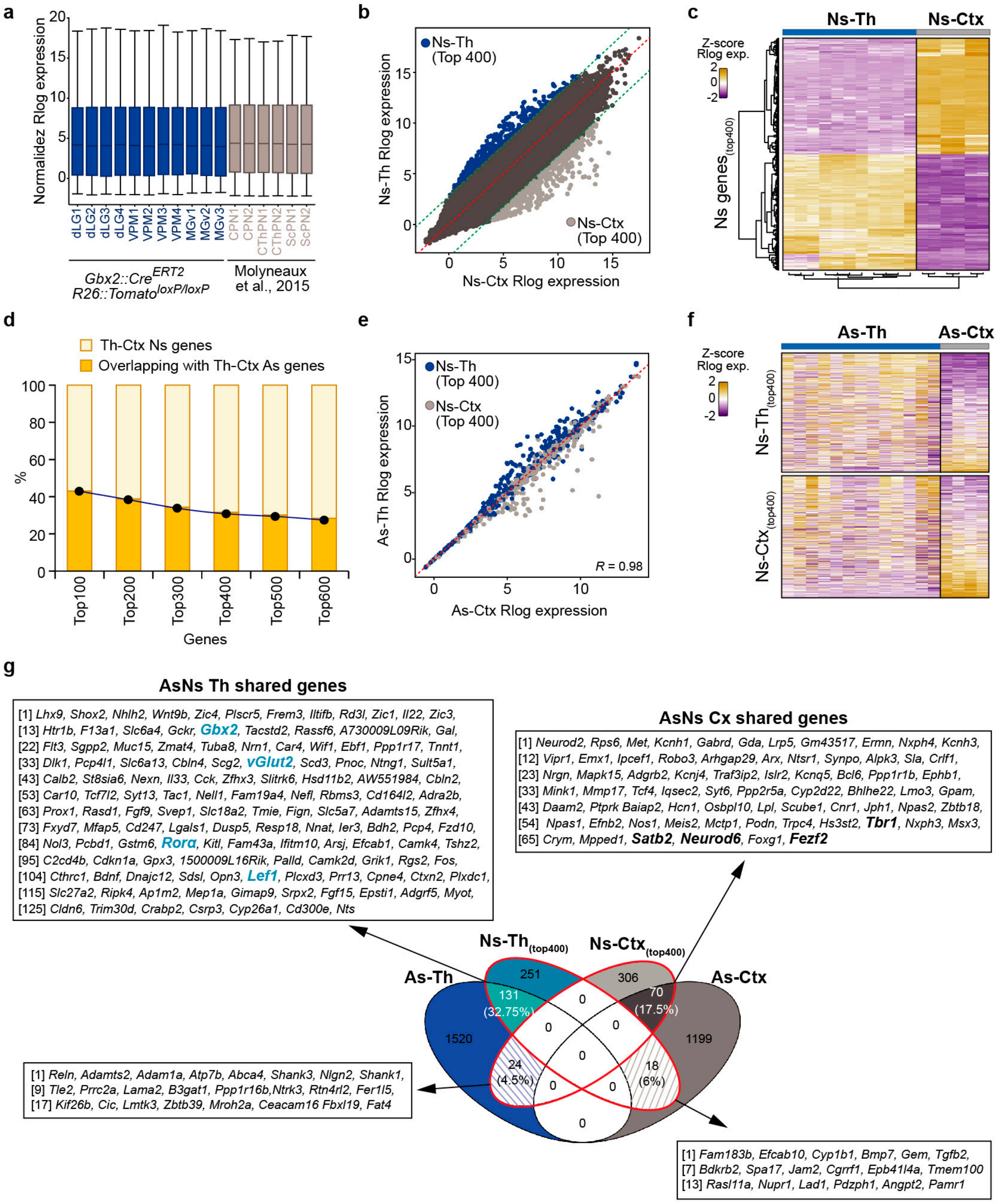


**Extended Data Fig. 2 | Neuronal genes shared in thalamic and cortical astrocytes. a,** Boxplot distribution of normalized Rlog expression of the transcriptome in thalamic neurons (Ns-Th, including dLG ($n = 4$), VPM ($n = 4$) and MGv ($n = 3$)) and cortical neurons (Ns-Ctx, including Callosal Projecting Neurons (CPN, $n = 2$); Corticothalamic Projecting Neurons (CThPN, $n = 2$); and Subcerebral Projecting Neurons (ScPN, $n = 2$)). **b,** Scatterplot of normalized Rlog expression showing the 400 top-ranked differentially enriched genes between thalamic and cortical neurons. **c,** Heatmap of Z-Score of normalized Rlog expression and unbiased clustering of 400 top-ranked differentially enriched genes between Ns-Th and Ns-Ctx samples. **d,** Percentage of overlapping of significantly DEGs between thalamic or cortical astrocytes and top-ranked DEGs in thalamic or cortical neurons according to the number of neuronal specific genes used for the analysis. **e,** Scatterplot of normalized Rlog expression in thalamic against cortical astrocytes for 400 top-ranked genes differentially enriched between thalamic and cortical neurons. **f,** Heatmap of Z-Scores of normalized Rlog expression in thalamic (As-Th) and cortical astrocytes (As-Ctx) for 400 top-ranked differentially enriched genes in thalamic and cortical neurons. In the heatmaps each row represents a gene, the columns are biological replicates and the color-code represents the normalized expression for upregulated genes in yellow versus downregulated genes in purple. **g,** Venn diagram showing shared genes between the Ns-Th and Ns-Ctx 400 top-ranked differentially enriched genes and the As-Th and As-Ctx significantly DEGs.

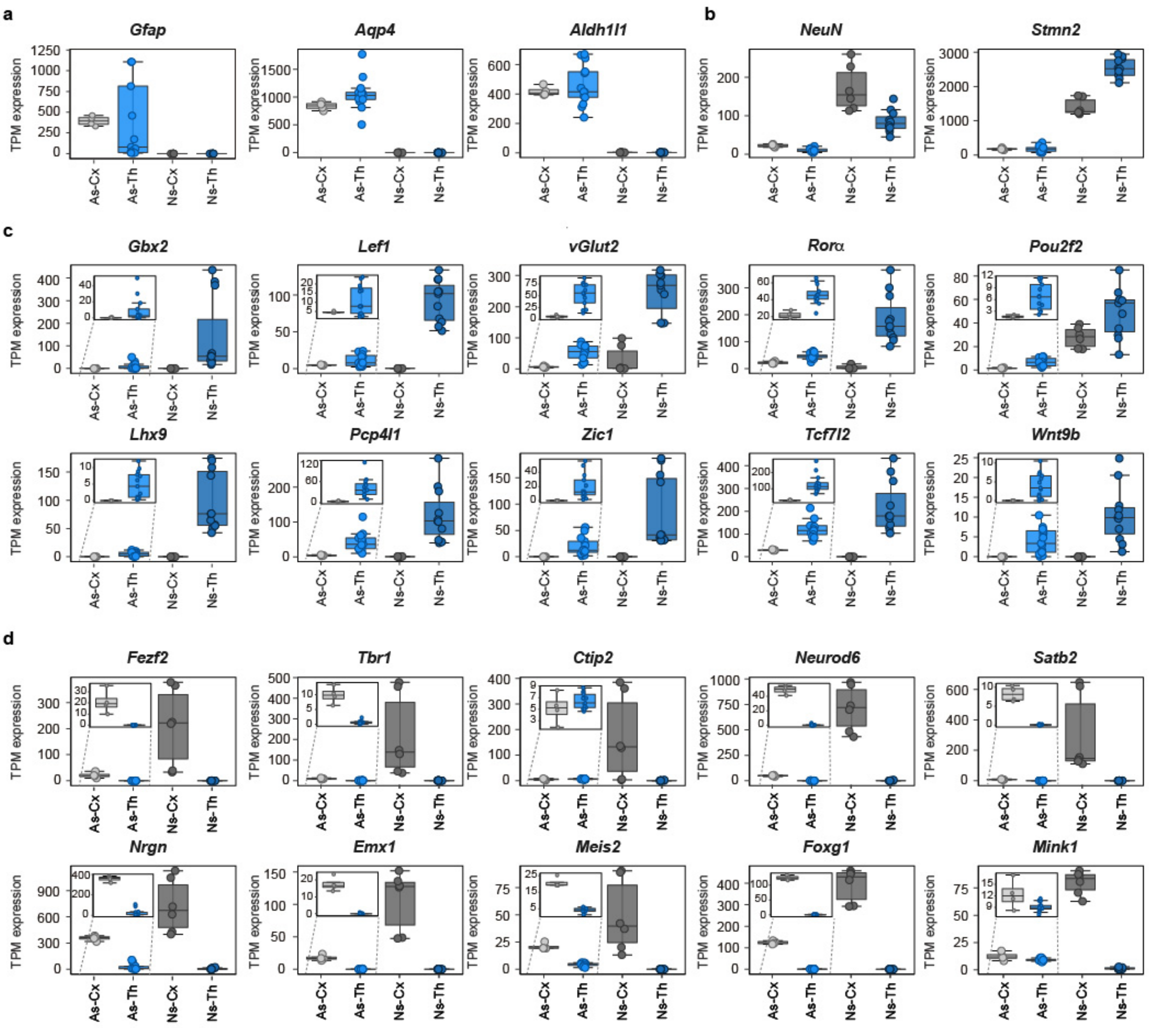


**Extended Data Fig. 3 | Comparison of expression levels of genes in neurons and astrocytes of the thalamus and cortex. a**, Box plots showing expression levels of pan-astrocyte markers *Gfap, Aqp4,* and *Aldh1l1* in cortical and thalamic astrocytes and neurons. **b,** Expression levels of pan-neuronal markers *Rbfox3 (*aka *NeuN)* and *Stmn2*. **c**, Expression levels of selected region-specific genes shared between neurons and astrocytes of the thalamus. **d**, Expression levels of selected region-specific genes shared between neurons and astrocytes of the cortex. As-Th, thalamic astrocytes; As-Cx, cortical astrocytes; Ns-Th, thalamic neurons; Ns-Cx, cortical neurons.

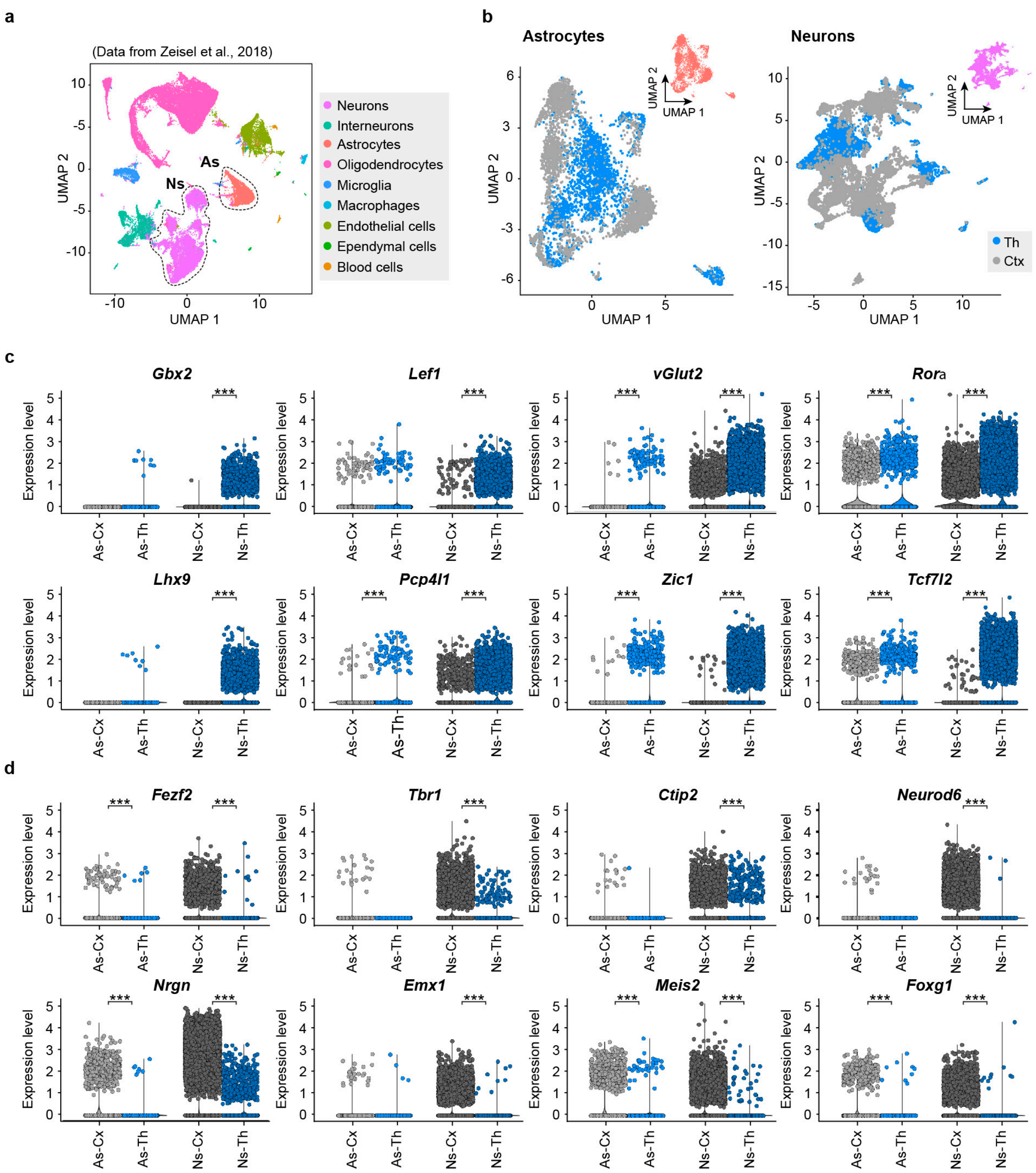


**Extended Data Fig. 4 | Single-cell sequencing analysis revealed region-specific genes between neurons and astrocytes.** **a,** UMAP plot showing the integration of single-cell RNA-seq datasets from mouse thalamus and cortex[18]. Astrocytic (As: 4763 cells) and neuronal (Ns: 15391 cells) populations are underlined. **b,** UMAP plots displaying differential clustering on astrocytic (blue dots correspond to 2131 thalamic cells and grey dots to 2632 cortical cells) and neuronal populations (blue dots correspond to 5856 thalamic cells and grey dots to 9535 cortical cells). The insets show the same clusterization but colored by cell type as in **a**. **c-d,** Violin and dot plots showing normalized expression levels in thalamic and cortical astrocytes and neurons of selected genes that are differentially expressed between thalamic and cortical neurons the bulk RNA-seq (As-Th, thalamic astrocytes; As-Cx, cortical astrocytes; Ns-Th, thalamic neurons; Ns-Cx, cortical neurons). * adj. $P$ value < 0.1; ** adj. $P$ value < 0.01; *** adj. $P$ value < 0.001.

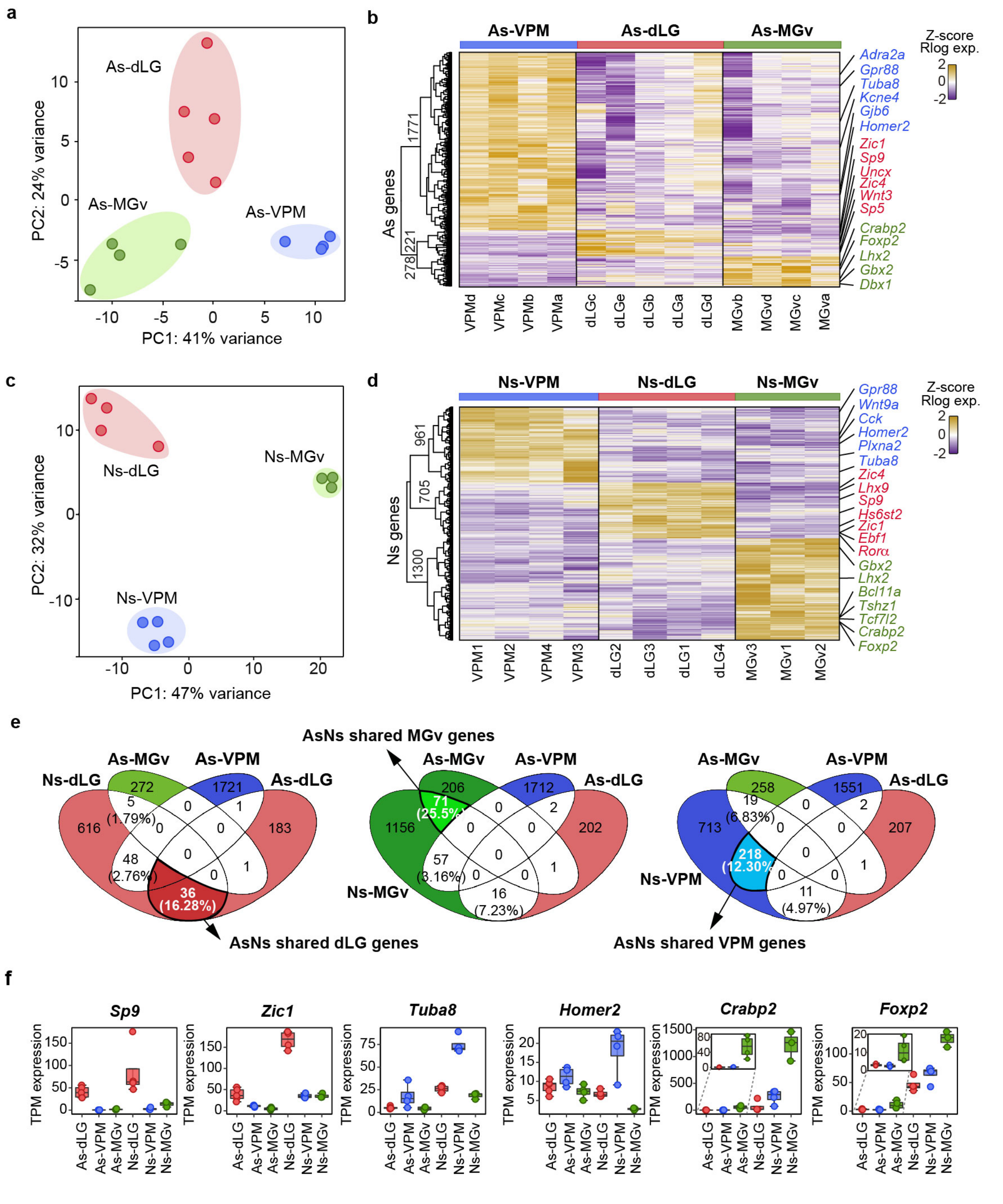


**Extended Data Fig. 5 | Thalamic nuclei-specific transcriptional signatures are shared in astrocytes and neurons. a,** Principal Component Analysis (PCA) of transcriptomes from astrocytes (As) isolated from the distinct sensory-modality thalamic nuclei (dLG ($n = 5$), VPM ($n = 4$) and MGv ($n = 4$)) at P7. **b,** Heatmap of Z-Scores of normalized Rlog expression and unbiased clustering of the significantly DEGs between astrocytes from distinct thalamic sensory nuclei. **c,** PCA of transcriptomes of neurons (Ns) from the distinct sensory-modality thalamic nuclei (dLG ($n$ = 4), VPM ($n$ = 4) and MGv ($n$ = 3)) at P0. **d,** Heatmap of Z-Scores of normalized Rlog expression and unbiased clustering of significantly DEGs between neurons from distinct sensory-modality thalamic nuclei. **e,** Venn diagram of significantly DEGs between astrocytes from distinct sensory-modality thalamic nuclei (As-dLG, As-VPM and As-MGv) and significantly DEGs between neurons from distinct sensory-modality thalamic nuclei (Ns-dLG, Ns-VPM and Ns-MGv). **f,** Box plots showing expression levels of nuclei-specific shared genes between astrocytes and neurons in the distinct sensory-modality thalamic nuclei. In the heatmaps, each column represents a biological replicate and the color-code represents the Z-score normalized expression for upregulated genes in yellow versus downregulated genes in purple.

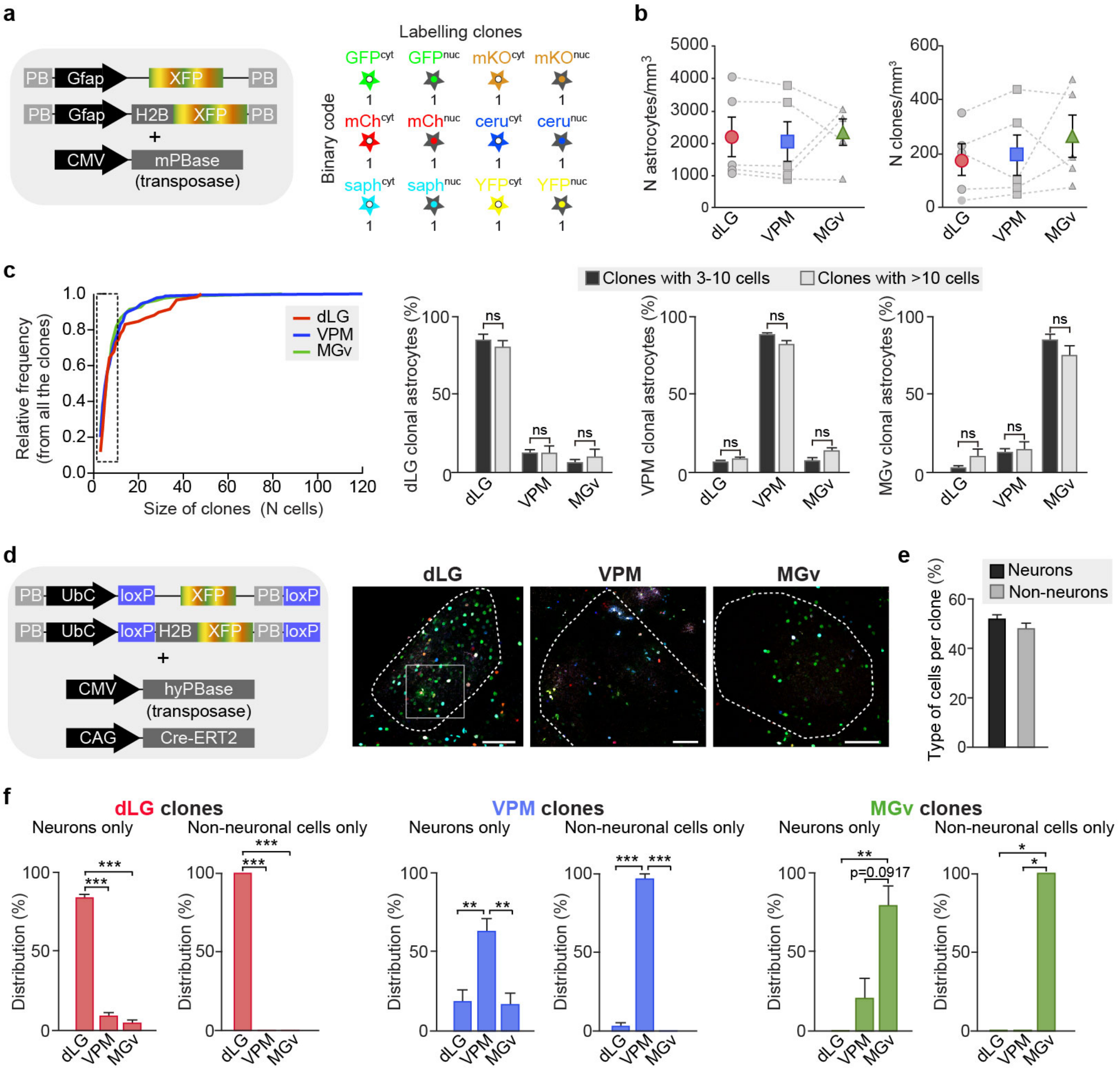


**Extended Data Fig. 6 | Astrocytes and neurons from the distinct thalamic nuclei are clonally related.** **a**, Left, Startrack plasmid combination used for the study of astrocytic clones. Right, binary code used for labelling the clones based on the presence and location of each fluorophore. **b**, Quantification of the distribution of the cells and clones through dLG, VPM and MGv nuclei in every electroporated animal ($n = 5$). Every point linked with a dashed line represents the data in each electroporated animal. **c**, Left, plot showing the relative frequency to the size of the astrocytic clones. Right, quantification of the distribution of the clones depending on their size ($n$ = 320 clones from 5 electroporated animals). **d,** Left, Startrack plasmid combination used for the study of clones including neurons and no-neuronal cells. Right, images showing electroporated cells in dLG, VPM and MGv of the same animal. **e**, Percentage of the presence of every cell type in the clones. **f**, Quantification of the distribution of the clones containing only neurons or non-neuronal cells across the three thalamic nuclei (*n* = 72 neuronal clones and *n* = 42 non-neuronal clones). Data are the means ± SEM. Scale bar, 100 μm. ns, not significant, $^{*}P < 0.05$; $^{**}P < 0.005$ and $^{***}P < 0.0005$.

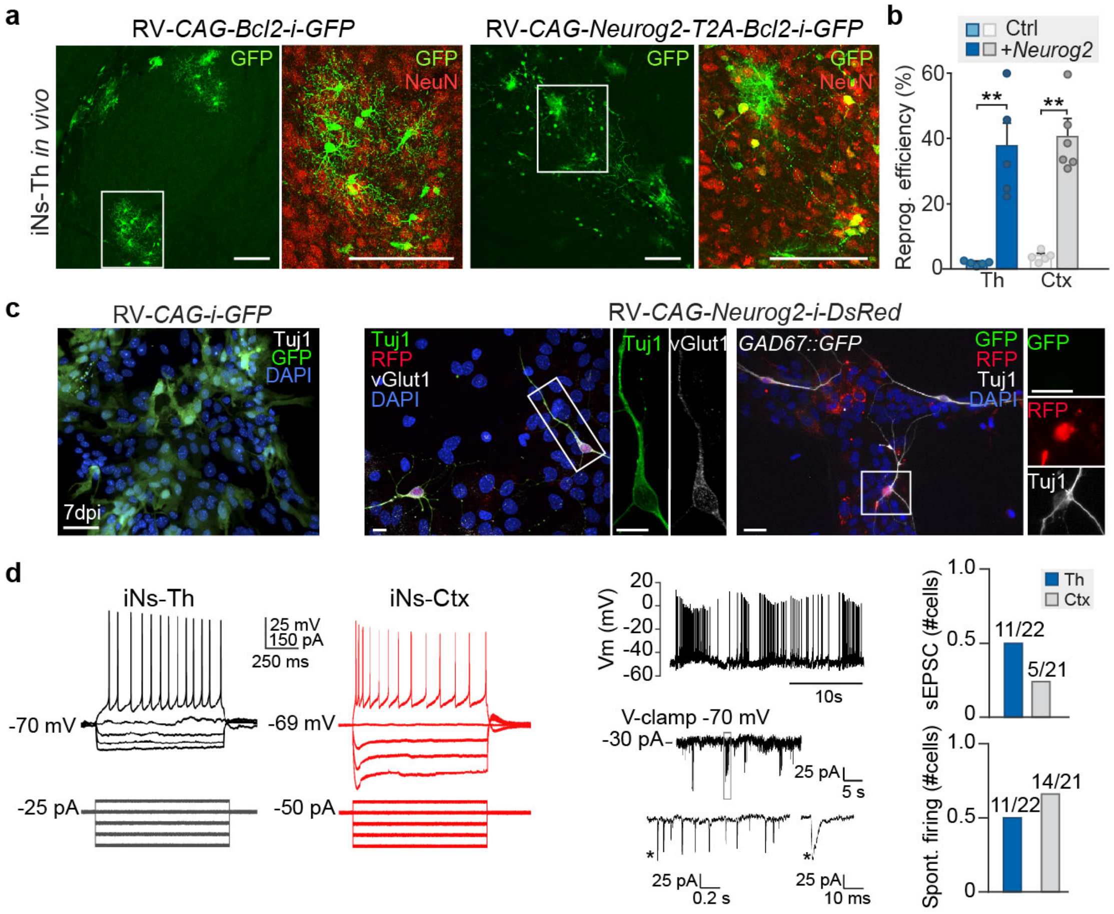


**Extended Data Fig. 7 | Thalamic astrocytes are reprogrammed into functional glutamatergic neurons by *Neurog2*. a,** Immunofluorescence of infected astrocytes *in vivo* with a control virus and a virus with Neurog2 and Bcl2 (*n* = 4-6 injected animals per condition). **b,** Quantification of the reprogramming efficiency *in vivo* in cortex and thalamus. **c,** Left, expression of Tuj1 in thalamic astrocytes infected with a control retrovirus containing *CAG-IRES-Gfp*. Right, immunostaining for vGlut1 and GFP in reprogrammed astrocytes (Tuj1+/RFP+) in thalamic cultures from wild type and *Gad67::Gfp* mouse after 10dpi. **d,** Left panel, whole-cell patch-clamp recordings in thalamic (left, black traces) and cortical (right, red traces) induced neurons (iNs). Central panel, whole-cell recording of the membrane potential (Vm) of a thalamic iN. Also, putative spontaneous excitatory postsynaptic currents recorded at -70 mV in a thalamic iN. Right graphs, number of recorded iNs from thalamic (*n* = 22 cells from 14 cultures; 28.3 ± 2.0 div) and cortical (*n* = 21 cells from 8 cultures; 30.8 ± 2.8 div) cultures that showed spontaneous excitatory post synaptic currents (sEPSC) or spontaneous firing. The data in **b** are the means ± SEM. Data in **b** contain the individual values for every injected animal or independent culture. Scale bar, 100 μm in **a**, and 50μm in **c** and **e**. *P < 0.05; **P < 0.005 and ***P < 0.0005.

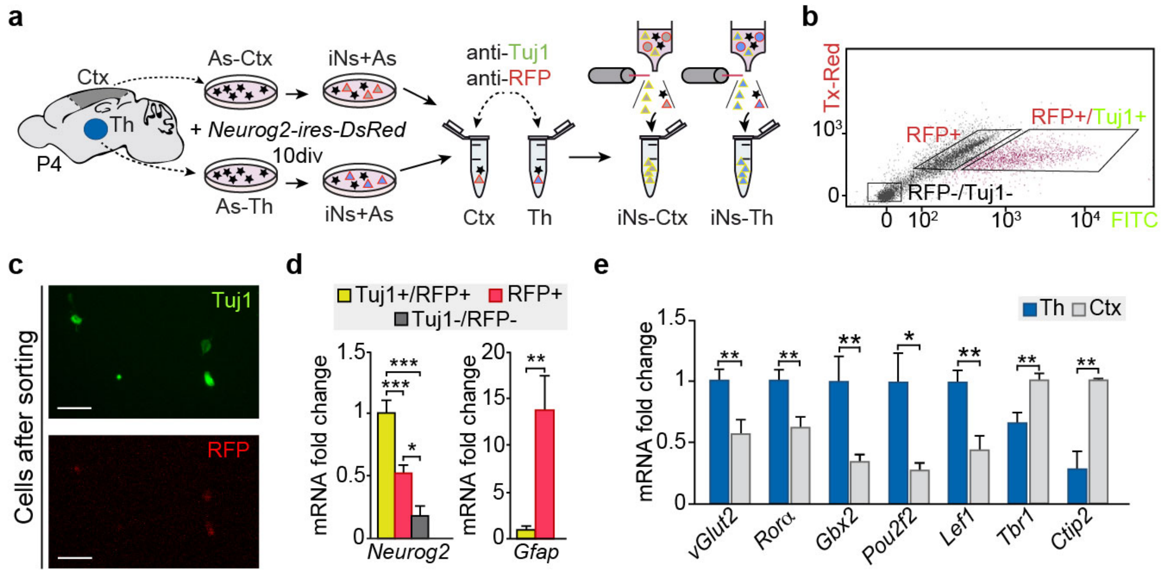


**Extended Data Fig. 8 | Region-specific genes expressed in reprogrammed neurons *in vitro*. a,** Experimental design for isolating induced neurons (iNs) with immunofluorescence followed by FACS sorting. **b,** Clouds of cells obtained by FACS sorting 10dpi after *Neurog2* viral induction. **c,** Double positive cells collected after sorting. **d,** Quantification of the expression of *Neurog2* and *Gfap* in the different sorted populations ($n$ = 8-12 independent cultures). **e,** Quantification of the differential expression of thalamic and cortical markers in the isolated iNs from thalamus and cortex ($n$ = 5-17 independent cultures). Data are means ± SEM. Scale bars, 50µm in **c**. *$P < 0.05$; **$P < 0.005$ and ***$P < 0.0005$

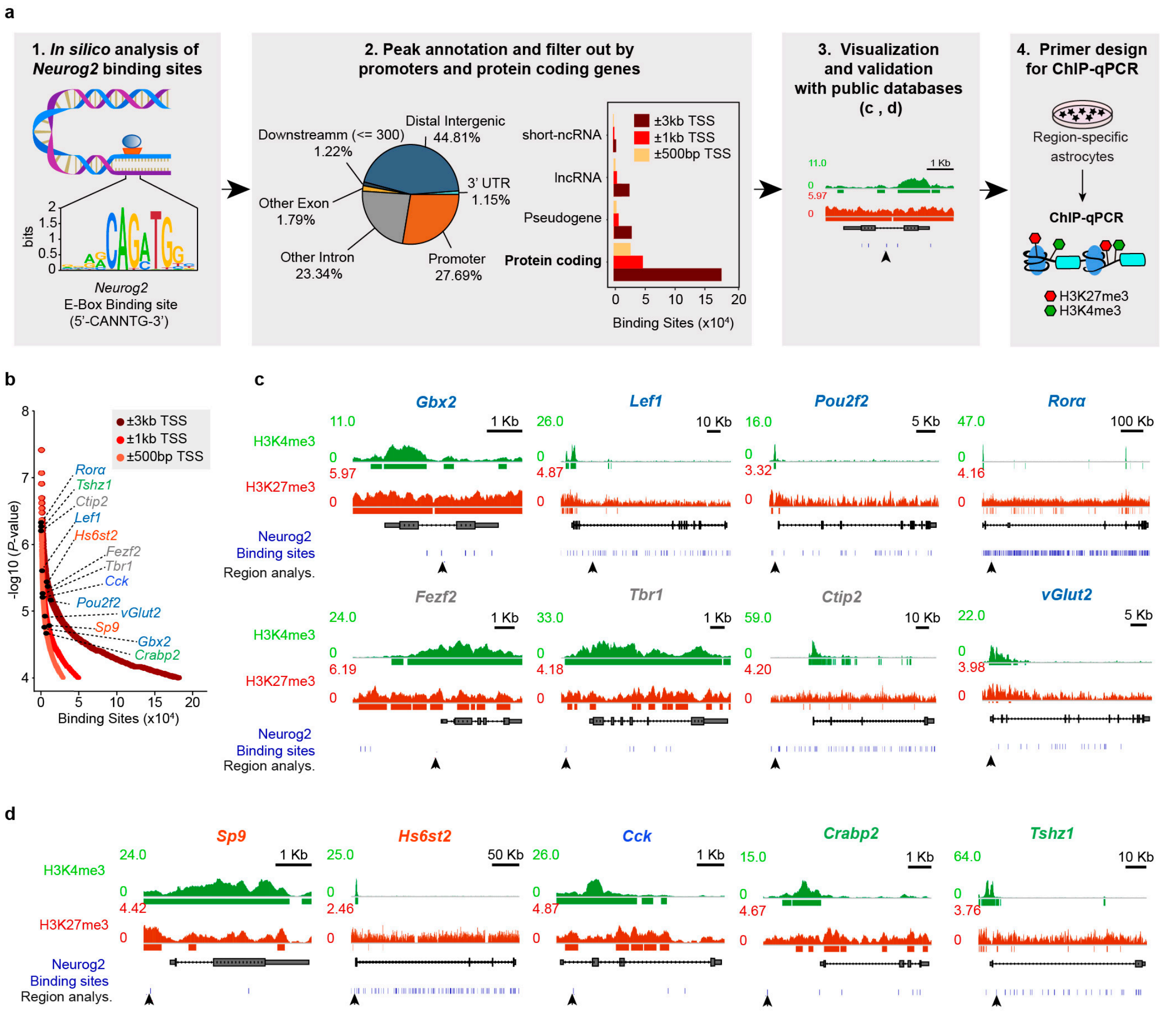


**Extended Data Fig. 9 | *In silico* analysis of candidate regions for the study of the epigenetic states of tissue-specific genes. a,** *In silico* pipeline analysis of Neurog2 targets binding sites used to design ChIP-qPCR primers. (1) Schema of the consensus binding site of Neurog2. (2) Peak annotation and filter out by promoter region and protein coding genes. Left, pie plot showing the genomic annotation by genic region. Right**,** number of binding sites of Neurog2 classified by gene biotypes extracted from promoter region and filtered out by genic proximity to transcription start site (TSS) (±3 Kb, ±1 Kb, ±500 bp from TSS). (3) Visualization and validation with public databases through genomic tracks. (4) Primer design for ChIP-qPCR. **b,** High-confidence putative binding sites of specific genes filtered by genic proximity (±3 Kb, ±1 Kb, ±500 bp) to TSS) found for Neurog2 across the whole genome by order of -log10 (*P*-value). **c-d,** *In silico* determination of selective promoter regions (black arrows) used to perform ChIP-qPCR analysis for thalamic, cortical and thalamic nuclei-specific genes, based on publicly available datasets of ChIP-seq from H3K4me3 and H3K27me3 histone marks and the putative binding sites for *Neurog2*.

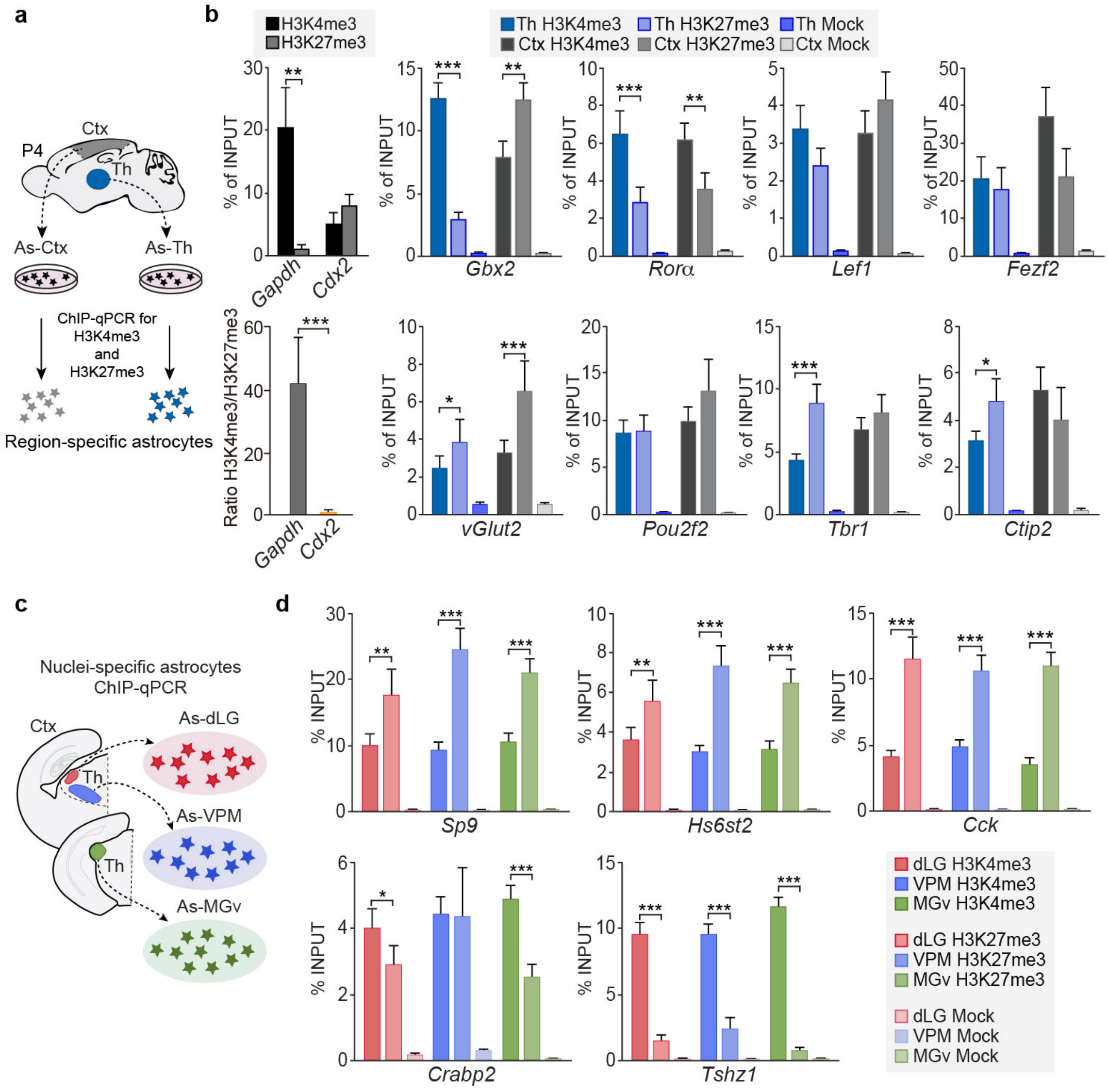


**Extended Data Fig. 10 | Region-specific genes show less epigenetic repression in the astrocytes in a tissue-dependent manner. a**, Experimental design in thalamic and cortical astrocytes. **b**, Quantification of the percentage of recovered input after the immunoprecipitation with H3K4me3, H3K27me3 and mock antibodies in thalamic and cortical cultured astrocytes ($n$ = 8-21 ChIP samples). **c,** Experimental design in astrocytes from dLG, VPM and MGv. **d,** Quantification of the percentage of recovered input after the immunoprecipitation with H3K4me3, H3K27me3 and mock antibodies in dLG, VPM and MGv cultured astrocytes ($n$ = 14-19 ChIP samples). Data are means ± SEM. $*P < 0.05$; $**P < 0.005$ and $***P < 0.0005$.

### Extended Data Tables

#### Extended Data Table 1 | Primary antibodies used for Immunohistochemistry.

| Antibody | Host | Concentration | Reference |
|---|---|---|---|
| **Anti-vGlut2** | Guinea Pig | 1:500 | Synaptic Systems #135404 |
| **Anti-GFP** | Chicken | 1:3000 | Aves Labs #GFP-1020 |
| **Anti-RFP** | Rat | 1:1000 | Chromotek #5F8 |
| **Anti-vGlut1** | Guinea Pig | 1:500 | Synaptic Systems #135304 |
| **Anti-NeuN** | Rabbit | 1:1000 | Millipore ABN78 |
| **Anti-Rora** | Goat | 1:200 | Santa Cruz sc-6062 |
| **Anti-Tbr1** | Rabbit | 1:500 | Abcam ab31940 |
| **Anti-Ctip2** | Rat | 1:500 | Abcam ab18465 |
| **Anti-Gfap** | Chicken | 1:1000 | Millipore AB5541 |
| **Anti-Dcx** | Rabbit | 1:500 | Abcam ab18723 |
| **Anti-Lef1** | Rabbit | 1:300 | Cell Signaling #2230 |

#### Extended Data Table 2 | Secondary antibodies used for Immunohistochemistry.

| Antibody | Fluorophore | Host | Concentration | Reference |
|---|---|---|---|---|
| **Anti-Rabbit** | Alexa 488 | Donkey | 1:500 | ThermoFisher #A21206 |
| **Anti-Mouse** | Alexa 488 | Donkey | 1:500 | ThermoFisher #A21202 |
| **Anti-Chicken** | Alexa 488 | Goat | 1:500 | ThermoFisher #A11039 |
| **Anti-Rat** | Alexa 594 | Donkey | 1:500 | ThermoFisher #A21209 |
| **Anti-Mouse** | Alexa 546 | Donkey | 1:500 | ThermoFisher #A10036 |
| **Anti-Chicken** | Alexa 546 | Goat | 1:500 | ThermoFisher #A11040 |
| **Anti-Rat** | Cy3 | Donkey | 1:500 | Jackson 712-165-153 |
| **Anti-Rat** | Alexa 647 | Goat | 1:500 | ThermoFisher #A21247 |
| **Anti-Mouse** | Alexa 647 | Donkey | 1:500 | ThermoFisher #A31571 |
| **Anti-Goat** | Dylight 649 | Rabbit | 1:250 | Jackson 305-497-003 |
| **Anti-Rabbit** | Alexa 647 | Donkey | 1:500 | Jackson 711-605-152 |
| **Anti-Chicken** | Alexa 647 | Goat | 1:500 | ThermoFisher #A21449 |
| **Anti-Guinea Pig** | Alexa 647 | Donkey | 1:500 | Jackson 706-605-148 |
| **Anti-Chicken** | Alexa 405 | Goat | 1:500 | Abcam ab175674 |

#### Extended Data Table 3 | Primers used for RNA expression levels.

| Target gene | Forward primer | Reverse primer |
|---|---|---|
| ***Gapdh* *** | CGGTGCTGAGTATGTCGTGGAGT | CGTGGTTCACACCCATCACAAA |
| ***vGlut2*** | TGCTACCTCACAGGAGAATGGA | GCGCACCTTCTTGCACAAAT |
| ***Rorα*** | TCCCACCTGGAAACCTGCCAGT | CCACGAGCGATCCGCTGACA |
| ***Tbr1*** | CAAGGGAGCATCAAACAACA | GTCCTCTGTGCCATCCTCAT |
| ***Ctip2*** | GGGCGATGCCAGAATAGAT | GGTAGCCTCCACATGGTCAG |
| ***Gbx2*** | GGCAACTTCGACAAAGCCGAGG | CCAGGCAAATTGTCATCTGAGC |
| ***Lef1*** | AACTCTGCGCCACCGATGAGATGAT | ATGACTTGATGTCGGCTAAGTCGC |
| ***Pou2f2*** | GCAAGAAGAGGACCAGCATC | GTGGTTCAAGAAGAGAGGCG |
| ***Fezf2*** | AACCCAAAAACTTCACCTGCG | TTGCACACAAACGGTCTAGC |
| ***Sp9*** | CGTCTATACTTGGGGAAGAGCC | GATCTTGTTGCAGGTCGCC |
| ***Hs6st2*** | CCAGGCTGAGACCTTCCAG | TCTGGACATGCCTCCATTCA |
| ***Cck*** | CCCTCAACTTAGCTGGACTGC | CTCTTCATGGCTTTGGATGGG |
| ***Crabp2*** | GGAATTTGAGGAGCAGACCG | TGCTCGCACACCATTTTGTT |
| ***Tshz1*** | ATGCCGAGGAGGAAGCAG | CGCAAATTCGCTCTCCTGAA |
| ***Gfap*** | AAGCTCCAAGATGAAACCAACCTGA | GCAAACTTAGACCGATACCACTC |
| ***Neurog2*** | AGAAGACCCGCAGGCTCAAG | CGTGGAGTTGGAGGATGACG |

* Housekeeping gene

#### Extended Data Table 4 | Primers used for ChIP-qPCR.

| Target gene | Forward primer | Reverse primer |
|---|---|---|
| ***Gapdh*** | GCCCCTTTCACCCACTGTAG | TCCCCTCACTGATCTCACCC |
| ***Cdx2*** | TAAGCTGTAGAGGCGGGGAT | ACACTCGTTAATCACGTAAGGC |
| ***vGlut2*** | TAGCTGATCTTTGCGAACGTG | TCCCCTAGGAAGTTGGCGAG |
| ***Rorα*** | CCTTTTTGTGAGCCTATTCTC | ATGGAACACAAACCACGA |
| ***Tbr1*** | AAATTGCTTCTACTTGATTCTTAG | CACTGAGCAGACTGCACCAC |
| ***Ctip2*** | GTCCTCCCTCATTGACCACA | CCCTCCGAACTGTAAACCAC |
| ***Gbx2*** | GTCCAGGACTGTTTGAAGACC | CACAAATGATATGTCTCAAGAAAGC |
| ***Lef1*** | CATGGTCCAGGTTATTCTCC | CTTAGCTTTATGATTCAGC |
| ***Pou2f2*** | GGATGGCCCTAGTGGATAGGA | TCACACAAAGGCGCAGGAC |
| ***Fezf2*** | AAAGTATCCCTTAGCCCTGCG | TCCATGCGCCACATCCTAATG |
| ***Sp9*** | CAGAGGCCAGAGCGACAATTA | CTGCTCCCACCGTACTCTTTT |
| ***Hs6st2*** | CCCCCTTCATGTTTGTGAGAC | CGCATGCACGAGAGCATTTC |
| ***Cck*** | CCAAAGCCCTGCTACACTTG | GGGTTTCTCTGCTGTTCGGA |
| ***Crabp2*** | CCCTGCCCCCTTGTTAGTTG | GGAGGGGGCAATATGCTAGG |
| ***Tshz1*** | TGTGCCGTCTCAATTTCCTC | AAGTTGGCAGGCTTGGTGTT |
